# The physiological costs of leadership in collective movements

**DOI:** 10.1101/2023.11.22.567987

**Authors:** Hanja B. Brandl, James A. Klarevas-Irby, Daniel Zuñiga, Christina Hansen Wheat, Charlotte Christensen, Fred Omengo, Cosmas Nzomo, Wismer Cherono, Brendah Nyaguthii, Damien R. Farine

**Author notes:** These authors contributed equally to this work and are joint first author (Hanja B. Brandl, James A. Klarevas-Irby). Senior author (Damien R. Farine).

## Abstract

Individuals can gain substantial benefits from collective actions^1–7^. However, collective behaviours introduce new challenges, like coordinating actions, maintaining cohesion, and meeting the needs of different individuals. When making collective movements, leaders are typically thought to gain disproportionate benefits through the choice of more beneficial resources^3^ and/or earlier access to resources^8^. However, reaping these benefits can also have costs. Being at the front of a group can increase physical exertion^4,9,10^ and predation risk^11,12^. Moreover, ending up in a leadership position (i.e., at the front), is a process of negotiation in many animal groups. Within-group differences in directional preferences are typically resolved by some individuals initiating directional movements, after which they are either followed (if they are successful in leading) or return to the group (if they fail)^13–30^. By combining data on movement initiations (using whole-group GPS tracking^31^ and individual heart rate from implanted ECG loggers) in wild vulturine guineafowl, we found significant increases in heart rate (and decreases in heart rate variability) during collective movements. We found that attempting—and failing—to initiate directional movement was particularly costly, with the highest costs when consensus among group members was low and when individuals acted against the majority. Increases in heart rate and decreases in its variability can indicate physiological stress, entailing increased energy expenditure and long-term physiological damage. These results suggest that behaviours often thought beneficial to individuals (by influencing group behaviours) are also physiologically costly, representing a constraint on group-living and explaining why sometimes individuals opt out of contributing to leadership.

## RESULTS & DISCUSSION

We studied the physiological consequences of collective movements and leadership in a group of wild vulturine guineafowl in Kenya. Vulturine guineafowl are large, predominately terrestrial birds endemic to East Africa. They live in multi-male multi-female groups with temporally stable membership^32^, ranging from 15 to over 65 individuals^33^. Despite steep dominance hierarchies^23,34^, groups reach consensus about where to go, maintaining cohesion, using shared decision-making^23^. Individuals move in their preferred direction (i.e. initiate) and are either followed (success) or not (failure) by others. All group members can lead^24^, but some engage in leadership more often than others. The highly dynamic nature of group movements (Movie S1) means all individuals occupy every spatial position (front, middle, back, side) within the group throughout the day.

To determine when and for whom, costs arise from individuals’ contributions to collective movements, we analysed data collected simultaneously from global positioning system (GPS) and electrocardiogram (ECG) loggers (Fig. 1; see materials and methods). GPS loggers were fitted to ∼70% of the members of a vulturine guineafowl group (N=20 of 28, collecting data at 1Hz, see^31^) from which we quantified individual spatial position, individual and group movement speeds, and both successful and unsuccessful movement initiations (following^24^). Additionally, an ECG logger was fitted to 65% of the GPS birds (N=13, from which we were able to collect high-quality data from 55% of the GPS birds, N=11), at 180Hz for 6s in every 20s window from 6am to 12pm, and corrected for individual daily resting heart rate. Data were collected on 31 days over 4.5 months (every fourth day), providing simultaneous information on group movements, individual spatial position, leadership, and heart rate.

**Fig. 1.**
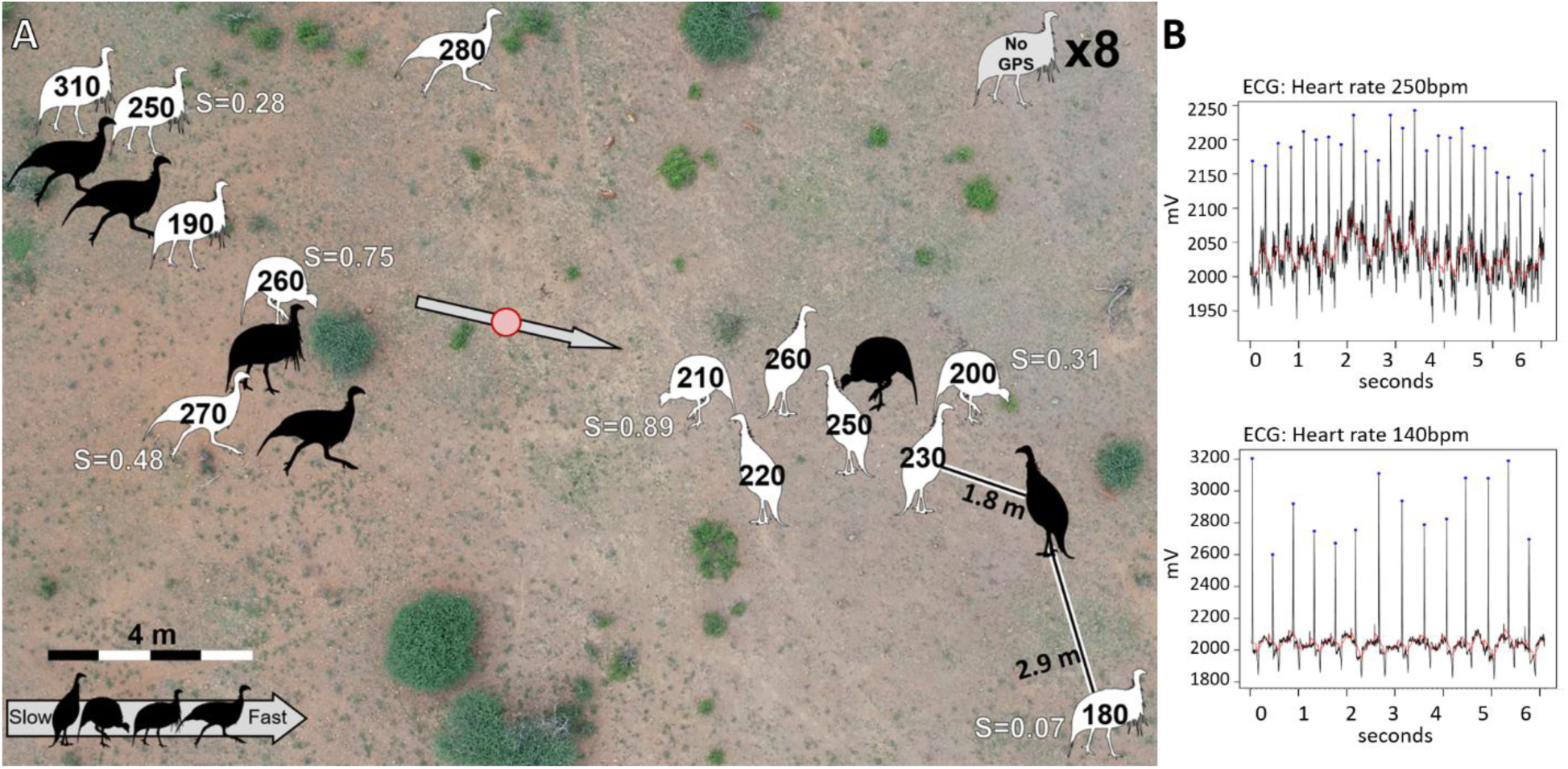
Snapshot of GPS and ECG data collected from a social group of vulturine guineafowl. (A) Spatial positions of individuals with only GPS loggers only (black) or both ECG and GPS loggers (white) from one ECG window. Black numbers within silhouettes indicate individual heart rates. Group centroid and movement direction are shown by the red dot and grey arrow, respectively. White text shows surroundedness (S) values for a subset of individuals. Nearest neighbour distances are given for the two leading individuals (black and white bars, bottom right). (B) ECG data from a 6-second window for an individual with high (top) and low (bottom) heart rates (both with low HRV). Background image courtesy of Blair Costello.

Heart rate, the frequency of heartbeats over time, is a reliable fine-scale proxy for autonomic nervous system activity and is modulated during physical exercise or in response to stressors^35,36^. Positively correlated with metabolic rate, increased heart rate also reliably indicates higher energy expenditure^37,38^. Guineafowl heart rates in our group ranged from 90 to 386 bpm, averaging at 181 bpm±31 SD, thus showing the potential for up to 4-fold increases in certain contexts. Consistently elevated heart rate can cause long-term physiological damage, such as oxidative stress^39^. In addition, heart rate variability (HRV), calculated as the root mean square of successive differences (RMSSD) between heartbeats^40,41^, serves as a physiological stress proxy. Reduced HRV, indicated by lower RMSSD values, describes the balance between sympathetic and parasympathetic activity and indicates active physiological stress responses.

### Moving as a collective increases the physiological costs of individual movement

To unravel how the activity of the group affects individual movement costs, we fit a linear mixed model (LMM, model 1, Table S1) to the baseline-corrected heart rate of vulturine guineafowl, based on their speed and position relative to the group. This revealed that collective movement strongly modulated individual heart rate (Fig. 2A). As expected, higher individual speeds corresponded to higher heart rates (model 1: coefficient(individual speed)±SE=22.97±0.78, t=29.36, p<0.001; Fig. 2A). However, the effect of individual speed on heart rate was contingent on group speed (model 1: coefficient(individual speed x group speed)±SE=13.97±0.97, t=14.41, p=<0.001; Fig. 2A). Specifically, the faster the group moved, the higher the individual’s heart rate for a given individual speed (Fig. 2A). A density plot showing the distribution of individual speeds in relative to different group speeds, with individual speeds being higher on average, is provided in Fig. S1. These results suggest that collective movements impose physiological costs over-and-above those incurred from moving at a given speed.

**Fig. 2.**
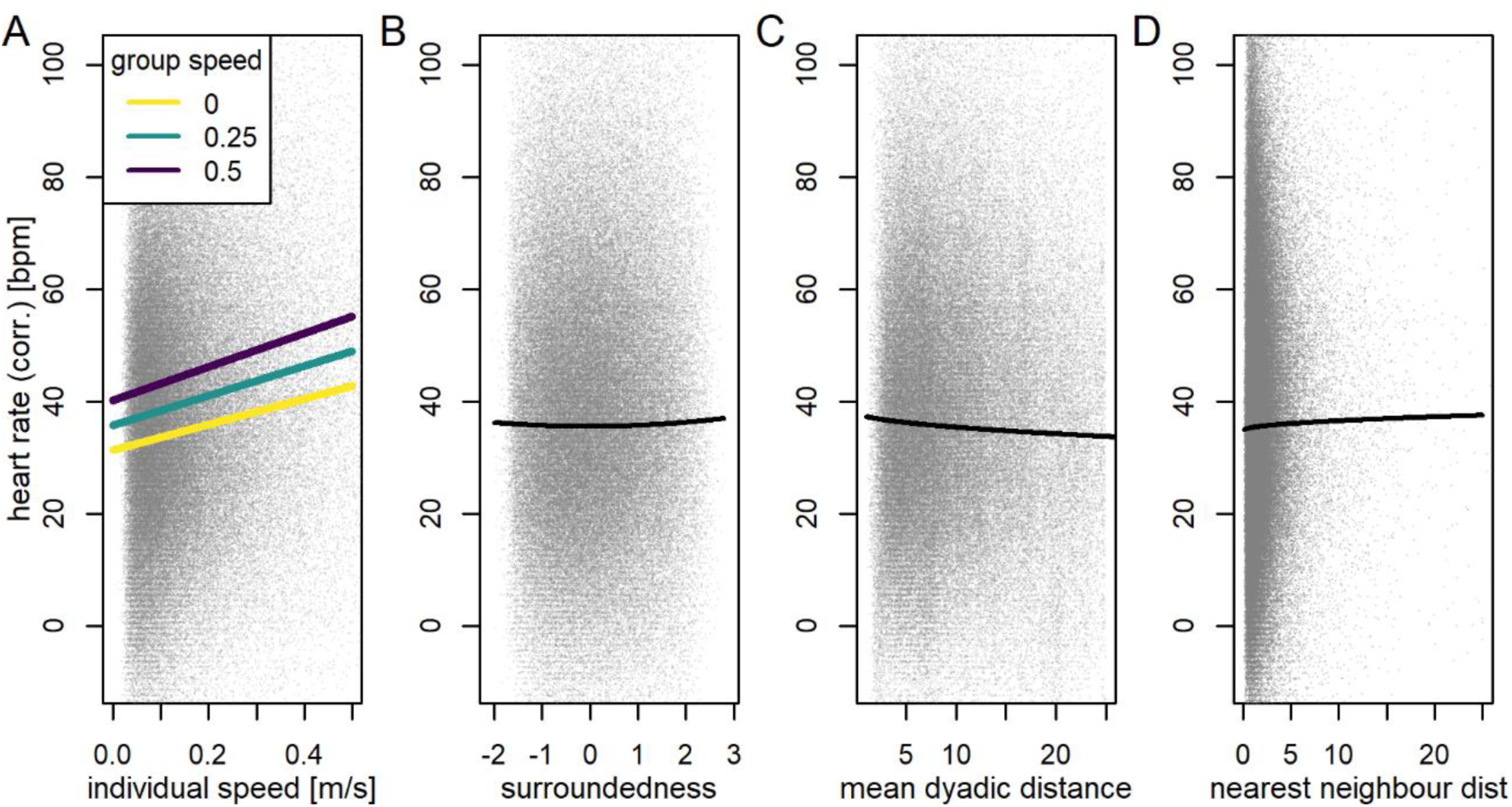
Heart rate responses to group movement and different within-group positions. Collective movement (A) increases heart rate, while (B) greater surroundedness, (C) higher group density, and (D) isolation from others only have minimal effects. Lines show model predictions from 118,553 ECG data windows. Corrected heart rates are the difference between individual absolute heart rate and their daily resting heart rate. All panels are from the same model (model 1, full results shown in Table S1), and average values are used for all covariates not shown: individual speed=0.1 m s^-1^ and group speed=0.1 m s^-1^ (representing slow movements), scaled and centred surroundedness=0 (range: −2.05–2.77), mean dyadic distance=9.8 (range: 1.13–25.0), nearest neighbour distance=2.11 m (range: 0.06–117.91). Grey points show 95% of raw data (excluding upper and lower 2.5%), jittered to reduce overlap.

### Spatial position has marginal physiological effects

We also analysed the contribution of individuals’ spatial position during group movements. Theory^42^ suggests individuals can gain safety from predators by positioning themselves between others to minimise exposure, while peripheral positions might increase danger and induce stress^43^. However, central positions—where individuals are typically more tightly packed—also requires individuals to pay substantially more attention to other group members (e.g. in scramble competition^44^), potentially counteracting potential safety benefits. Our data show that individuals pay only marginally greater costs—from a physiological perspective—when they are more central or more peripheral (Fig. 2B–D). Specifically, heart rate was slightly higher at both extremes of surroundedness, a robust measure of spatial centrality^31,45,46^ (model 1: coefficient(scaled surroundedness^2^)±SE=0.18±0.07, t=2.41, p=0.016; Fig. 2B and Table S1). Correspondingly, heart rate also increased slightly when groups were more tightly packed (model 1: coefficient (sqrt(mean dyadic distance))±SE=-0.89±0.10, t=-9.3, p<0.001; Fig. 2C) and when individuals were more isolated (model 1: coefficient (sqrt(nearest neighbour distance))±SE=0.54±0.15, t=3.61, p<0.001; Fig. 2D). Increases in heart rate corresponded to decreases in HRV (model 4, Table S2, Fig. S2), consistent with stress arousal. However, while the direction of all spatial positioning effects aligned with theory, changes in heart rate and HRV were small—at least one order of magnitude lower than movement speed effects—and thus unlikely to impose substantial costs. Further, because spatial positions are highly dynamic in moving animal groups^9,28,47^ these effects are unlikely to accumulate into significant physiological burdens.

Overall, we found clear evidence for collective movements driving increases in individual physiological costs, but only marginal effects of the moment-by-moment spatial position of individuals—relative to group members—on these costs. Previous studies link individual physiological state to spatial position (e.g., unfed individuals can be found at the front of groups^9,23^, a.k.a. leadership according to need^48^). Such patterns reflect differences in preferences among group members, creating potential conflict within the group over movement decisions. A key question is therefore whether these conflicts—and the role of leadership in resolving them— introduces physiological costs at the individual level.

### Disagreement drives a physiological cost of leadership

Central to functioning as a social group is the ability to overcome difference in preferences among group members, as failure to reach consensus can lead to group splits^49,50^. In many group-living animals, including primates^28^, other mammals^51^, fish^52^, and birds^23^, group members reach consensus about movement directions through a process akin to voting. Some group members initiate movement by moving away from others, in their desired direction of travel, and conflicts in preferences—when more than one directional preference exists—are resolved by selecting the direction of the majority of initiators^53^. While leading can offer benefits, such as access to preferred resources, almost nothing is known about the costs that leadership entails, particularly when attempting to lead a group with a degree of internal conflict.

To test the prediction that directional disagreement among group members (i.e. leadership conflicts) can be costly, we summarised co-occurring initiations detected from the GPS data into events and identified the level of directional agreement among co-initiators according to the distribution of their movement vectors (using the methods from^24^). Initiations are inferred from dyadic changes in distance (per^28^), such that an initiator is the individual responsible for a temporary increase in dyadic distance with the other individual becoming a potential follower (i.e. it may follow, causing the initiator to be successful, or not, causing the initiator to be ‘anchored’ by returning). This prediction was supported by an LMM (model 2, Table S3) fit to the corrected heart rate, together with group speed, directional agreement among simultaneous initiators (potentially) acting on the same followers, and the number of potential followers (i.e., the number of group members that an initiator simultaneously increased its dyadic distance with) as a measure of behavioural disagreement, alongside other movement and position variables (from model 1). We first found that, when initiating movement, individuals had the highest heart rate when the overall directional agreement among initiators was low and when there were many potential followers (suggesting that the initiator was acting against a majority, and a stronger consensus among group members). This effect was also strongest when the group was on the move (model 2: coefficient(directional agreement x group speed x N potential followers)±SE=-24.77±9.87, t=-2.51, p=0.012; Fig. 3A-B; see Table S3 for individual effects and other interactions). We also found the same pattern (low agreement with high numbers of followers) being linked to higher stress levels (i.e., lower HRV; see model 5, Fig. S3, Table S4). These results confirm that initiations are costly and that these costs are highest when disagreements arise during collective movements.

**Fig. 3.**
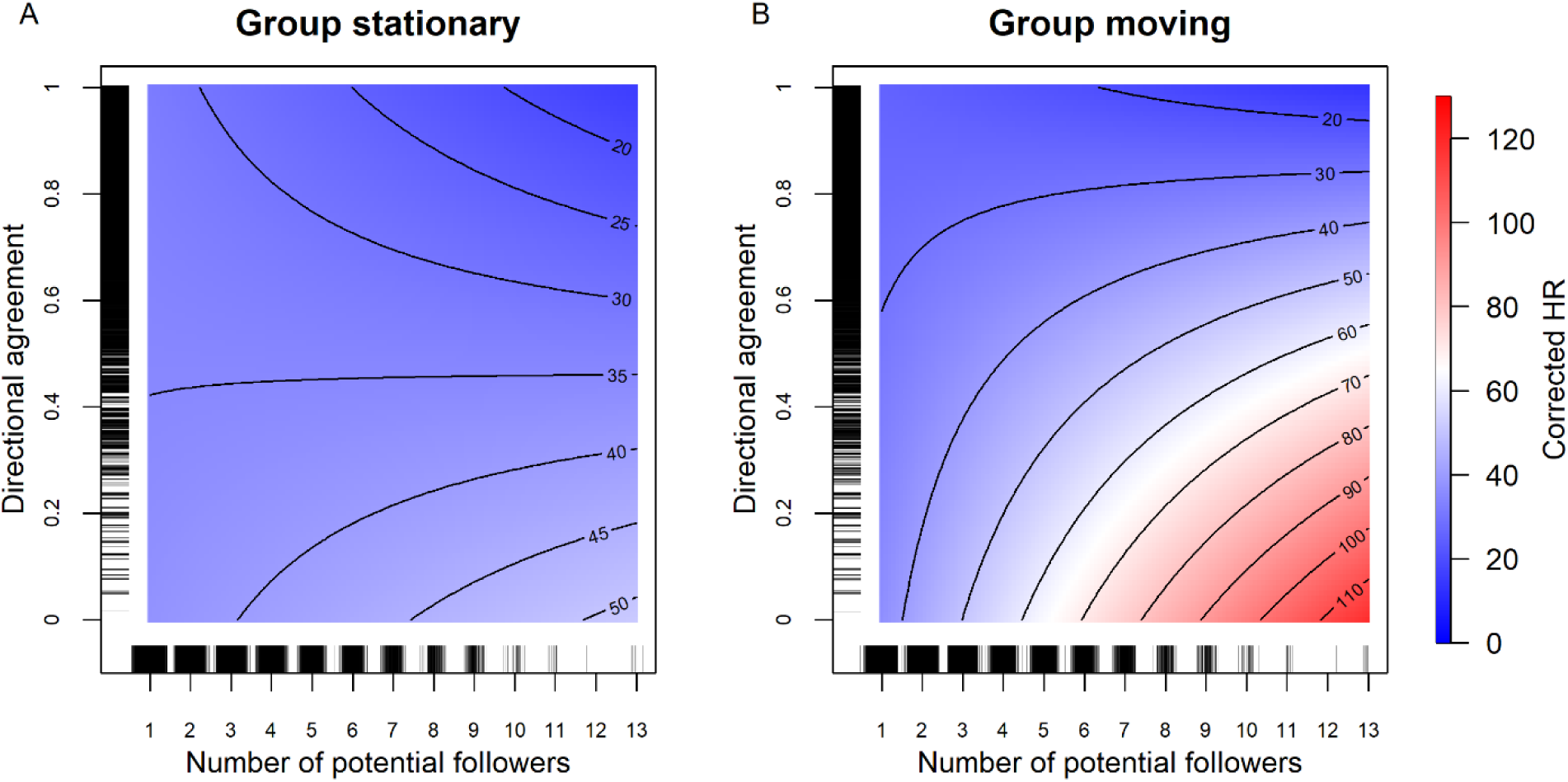
The cost of leadership is greatest with more potential followers and low directional agreement among initiators, especially when the group is moving. Model predictions from 64,126 ECG data windows (20 s each) in which the focal individual was initiating movement. The color-coded heart rate measure (HR) is corrected and thus shows the increase in heart rate, in beats per minute, relative to their resting heart rate. All panels are from the same model (model 2, Table S2), with group speed set to either 0 m s^-1^ “not moving” (A) or 0.2 m s^-1^ “moving” (B), and all other terms set to 0 as these only had weak effects. Density distributions of raw data points are shown along the axes.

### Failed initiations are costly, especially when directional disagreement is high

While initiating movement is costly, a final question is whether the outcome of the initiation modulates these costs. We determined the outcome of initiations (successful if the initiator was followed, unsuccessful if not) by investigating the period after the initiation, predicting that failed initiations would be more costly. An LMM (model 3, Table S5) fitted to the post-initiation outcomes, with an interaction between the outcome (individual initiated successfully or not) and the level of directional agreement among initiators prior to the event confirmed our prediction. Individuals had the highest heart rates (controlling for spatial position and movement) when their initiation failed, especially when agreement among the initiators was low (model 3: coefficient(successful x directional agreement)±SE=7.23±3.29, t=2.20, p=0.028; Fig. 4; see Table S5 for each individual effect). These results were not simply explained by movement, as individuals that were followed typically moved faster than those that were unsuccessful during the same period (Fig. S4). Thus, these results confirm that the social context in which an individual initiates movement, and the collective outcome of that initiation, can have physiological consequences.

**Fig. 4.**
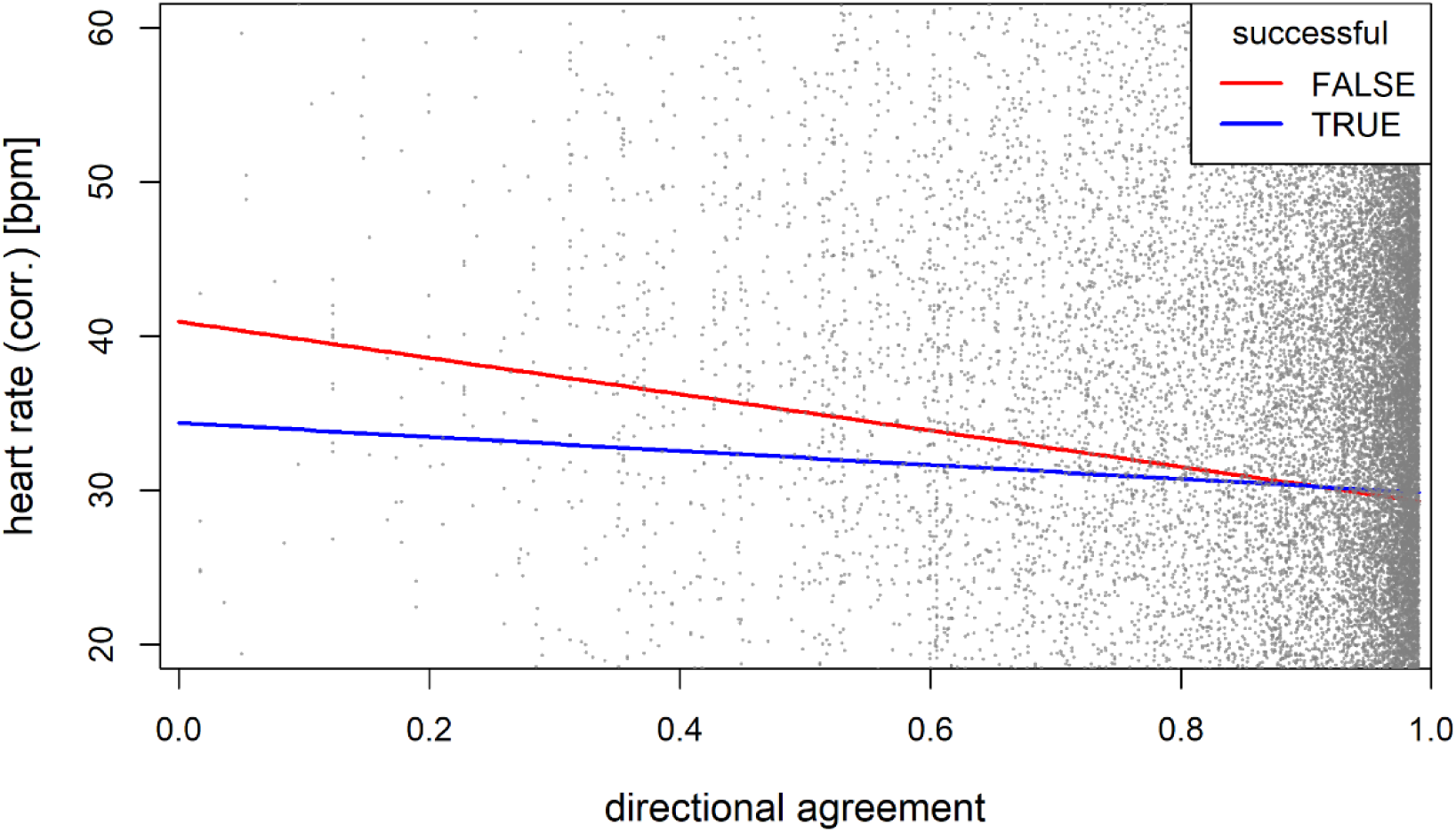
Unsuccessful initiators have a higher heart rate than successful initiators, especially in presence of directional conflict among initiators (low agreement). Model predictions from 22,398 ECG data windows (20 s each) where an individual was either successful (TRUE; blue line) or unsuccessful (FALSE; red line) in initiating. Corrected heart rate reflects increases relative to individual daily resting heart rate. in beats per minute. Fits are from model 3 (Table S5), with other variables set to 0 as these only scaled the results up or down along the y-axis. Grey points show 95% of the raw data. We found little evidence that initiation outcomes impacted HRV (Table S6, Fig. S5).

## Conclusion

While group living is an advantageous strategy for individuals across species, we have demonstrated that the dynamics of collective behaviour can also have major physiological consequences. Specifically, we found that collective actions drive increased heart rates over-and-above individual movement and spatial positioning alone. For example, an individual’s heart rate at a given movement speed is higher when moving as part of a collective (i.e. when other group members are also moving vs. when the group is stationary), likely contributing to additional energy expenditure over the course of the day. Conversely, the physiological cost associated with peripheral positions in the group—which in theory increases exposure to predators—is relatively negligible. The physiological costs of collective actions are most evident in individuals attempting to lead when there is directional conflict among group members. Initiators have the highest heart rates i) when the directional agreement among initiators is low, ii) when they are trying to influence more group members (i.e. initiating against the majority), and iii) when the group is already on the move (and accounting for other factors such as individual speed). Moreover, the outcome of initiations, and the consequences that these have on individuals, is also affected by conflict within the group. Failed initiators exhibit higher heart rates with low directional agreement among initiators, suggesting that the highest costs are borne by unsuccessful leaders.

While we focused predominately on heart rate, the effects of collective movements and initiations against directional and numerical conflicts also corresponded to lower HRV. While increased heart rate is linked to energy expenditure, lower HRV suggests that these situations also result in increased physiological stress. Surprisingly, we did not find evidence that failed initiation attempts also induce stress responses. However, the results on HRV might not be as robust as the heart rate measure, as calculating RMSSD from very short time windows (6s of data) makes it more sensitive to irregularities, thereby weakening the strength of the signal in the data.

While our study was conducted in only one species, we expect the results to reflect general patterns across group-living species. Several factors suggest that the effects observed in our study are relatively weak compared to what may be possible in other social groups and/or other species. For example, for logistical reasons we studied a relatively small group (N=28, see materials and methods for details), whereas vulturine guineafowl typically form groups of up to 65, sometimes over 90 individuals—conditions likely to exacerbate the challenge of making collective decisions and to amplify directional and numerical conflicts. Further, vulturine guineafowl exhibit higher levels of collective agreement during collective movements^56^, and lower rates of agonistic interactions^34^, than other species that make similar collective movements in similar environments (e.g. olive baboons, *Papio Anubis*)^28^. Thus, the cost of being more surrounded, initiating, or failing to lead may be substantially greater in primates and other mammals that live in cohesive groups than what we observed in vulturine guineafowl.

Overall, our results help to explain why individuals express a range of different social strategies within social groups over time, sometimes acting as leaders to secure benefits, and sometimes acting as followers to avoid costs. Ultimately, the asymmetric benefits of group living are accompanied with asymmetric costs for individuals and social interactions can introduce proximate (physiological) constraints shaping evolutionary trade-offs.

## ACKNOWLEDGMENTS

We thank Maureen Kamau and Edel Odhiambo for assistance during the implantation procedures, the Mpala Research Center for logistical support, and the Vulturine Guineafowl Research Programme field team—Janet Kariuki, John Wanjala, Kennedy Sikenykeny, Mary Ngugi, and Monicah Wambui—for their help with data collection and data downloads. This research was funded by the European Research Council (ERC) under the European Union’s Horizon 2020 research and innovation programme (grant agreement number 850859), an Eccellenza Professorship Grant of the Swiss National Science Foundation (grant number PCEFP3_187058), the Deutsche Forschungsgemeinschaft Centre of Excellence 2117 “Centre for the Advanced Study of Collective Behaviour” (ID 422037984), the Max Planck Society, and the Swedish Research Council (grant number 2019-06407).

## AUTHOR CONTRIBUTIONS

Conceptualization: DRF

Methodology: HBB, JKI, DZ, CHW, CC, FO, CN, WC, BN, DRF

Investigation: HBB, JKI, CC, DRF

Visualization: HBB, JKI, CC, DRF

Funding acquisition: DRF, CHW

Project administration: FO, CN, DRF, CHW

Supervision: DRF

Writing – original draft: HBB, DRF

Writing – review & editing: HBB, JKI, DZ, CHW, CC, FO, CN, WC, BN, DRF

## RESOURCE AVAILABILITY

### Lead contact

Further information and requests for resources and reagents should be directed to and will be fulfilled by the lead contact, Hanja Brandl.

### Materials availability

This study did not generate new unique reagents.

### Data and code availability

All data and code will be made publicly available as of the date of publication.

## SUPPLEMENTARY MATERIALS AND METHODS

### Materials and Methods

#### Study system and selection of social group

We conducted our study on wild vulturine guineafowl at the Mpala Research Centre in Laikipia county, Kenya (0.292 N, 36.898 E). The environment at our study site consists of mixed acacia bushland, with dense scrub interspersed with dirt roads and open glades, with seasons defined by rainfall (typically two wet and two dry seasons per year). Vulturine guineafowl are active during daylight hours (ca. 06:00-19:00), moving between dense brush and glades, which is where the majority of foraging and observable social interactions take place, and roosting in stands of acacia trees in the evening. This study took place from February-July 2021, with the early parts of the study taking place under intermediate seasonal conditions and slowly drying out due to the onset of what would eventually become a severe drought in 2022 and early 2023.

Several factors make vulturine guineafowl a good model for such a study. First, they are large, and their body weight (1.3 – 1.8 kg) means that they can carry equipment capable of collecting high resolution data over several months. Second, they are almost exclusively terrestrial, flying only to cross rivers or to go into trees (e.g. to roost in the evening), or when making short-distance, rapid escapes during predator attacks. Third, they are seasonal breeders meaning that by studying outside of this period we avoided any confounds arising from reproductive behaviours. Fourth, groups are extremely cohesive, often touching each other while making collective movements (see Movie S1). Previous work on the system has also demonstrated that their society is relatively egalitarian [S1].

We had initially aimed to collect data from two social groups, our reported group (N=28) and a second, similar-sized group (N=29). However, delays in permitting and worsening drought caused the second group to merge with another, much larger group, and when trapping we only captured half of that group’s members. Hence, we could not collect sufficient data from the second group to include them, and our strategy of targeting two groups meant that our deployments were limited to smaller groups. Because these two groups were the smallest in our population at the time, we expect that our results are relatively conservative.

#### ECG logger implantation and data collection

ECG loggers (ECG-tag 1AA2, e-obs GmbH, Grünwald, Germany) were sterilized using ethylene oxide and subsequently implanted into the thoraco-abdominal cavity of 6 males and 7 females between February 25 and March 2 2021. Birds were anesthetized with isoflurane 5% and oxygen (1-3 L/min); butorphanol 1.5 mg/ml was injected in the pectoral muscle to provide analgesia during surgery and ringer solution (20 ml/kg, SC) was injected for fluid maintenance. An incision was made in the linea alba (∼2 cm length) where the ECG loggers were implanted. The longer electrode was placed close to the heart and after confirming visually (QRS complex) that the live ECG signal was of good quality, the logger was fixed to the abdominal wall using an absorbable suture (Monosyn® 4/0, B. Braun AG, Melsungen, Germany). Post-operative pain was managed with meloxicam IM 0.5 mg/kg and Marblofloxacin 5 mg/kg IM was injected as infection prophylaxis. Loggers weighed 25 g or about 2% of the body weight of the smallest vulturine guineafowl tagged (1.3 kg). We used data starting from March 10 2021 onward, to give individuals time to recover from the implantation.

The ECG loggers were programmed to record one 180 Hz burst of data for 6 seconds every 20 s. The raw data was stored on the battery-powered devices and was downloaded via radio antennas together with the GPS data. The sampling strategy was designed to maximise the longevity of the battery life of the ECG loggers while still collecting data for the majority of the sampling period. ECG tags were programmed to collect data from 6 am until 12 pm, as the activity of birds starts to decrease substantially after 10 am [S2]. We did not include the first 30mins of each day into the analyses, because this is the period where individuals come down from their nightly roosts, which was accompanied with higher heart rates than during the rest of the day, and thus did not seem representative of their ‘normal’ behaviour. Because of clock drift, tags started collecting data progressively earlier over the course of the study, allowing us to use the pre-6 am data to extract the daily resting (baseline) heart rate of each individual.

#### ECG-based measures

The ECG data was analysed with a custom-made peak detection algorithm designed to detect the R-peaks from the ECG signals, in the R environment [S3]. We automatically filtered the data for artifacts, removing data windows with unusual gaps or anomalous R-R intervals or RMSSD (Root Mean Square of Successive Differences) values. After the automated error checking procedure, we also visually inspected 20 random ECG windows per individual per day to ensure correct R peak identification. In the quality control pipeline, we removed 8,2% of originally 206,835 data windows. We then calculated the heart rate in beats per minute from the 6 seconds of collected data windows every 20 seconds, and subtracted the resting heart rate, which ranged between 111 and 165 bpm, for each individual to obtain a measure of heart rate increase relative to their resting heart rate. We also obtained the root mean square of successive differences (RMSSD) by first calculating each successive time difference between heart beats (R-peaks) in ms, squaring each of these values, and taking the square root of their mean as a measure of heart rate variability. RMSSD is a commonly used proxy for physiological stress, particularly in short-term measurements [S4]. All of these decisions were made, and implemented, prior to analysing the data for our research questions.

#### GPS data collection

We fitted solar-powered GPS loggers (e-obs 15 g solar) to the majority of vulturine guineafowl in a social group. Tags were fitted using a backpack harness using Teflon ribbon and elevated using several layers of rubber matting. The total weight of the GPS logger plus backpack and harness is less than 2% of the body weight of all individuals (mean adult weights: females 1410 g, males 1625 g). For birds individuals fit with both GPS and ECG loggers, combined weight of all materials (ECG, GPS, harness, and rings) was less than 4% body weight (mean 3.3%, range 2.8-3.8%). GPS loggers were programmed to collect data every fourth day (the same day for all tags), ensuring that loggers had sufficient time to fully recharge batteries, allowing us to collect 1 Hz (one GPS location per second) data for the majority of the day (from 6am until the battery depleted, which was always later than the ECG loggers switched off, see below). Data were downloaded every second night using a VHF antenna, as part of a long-term study in this population.

While we were unable to trap all group members, the coverage (∼70% of group members) was—over the course of the study—comparable to previous studies. For example, the study by Strandburg-Peshkin et al. [S5] on leadership in olive baboons (*Papio anubis*) was based on between 50% and 80% of group members being fitted with a GPS collar. Our prior testing suggests that estimations of relative positions are highly accurate with these tags, even when not all individuals are sampled in the group (e.g. median error in surroundedness is ∼0.1 when 70% of the group is tracked and after accounting for GPS error [S6]).

#### GPS-based measures

For each second of each day of logging, i.e. from 6am (birds leave the roost around 6:15am) until 7pm (birds enter the roost around 6:50pm), we extracted a range of group- and individual-level metrics from all GPS-tagged individuals (i.e., including those not implanted with an ECG logger). At the group level, we first extracted the position of the group centroid (mean of all individual positions) as a proxy for the group’s location. We then calculated the speed (in meters per second) and bearing of the group movements from the displacement of the group centroid. Finally, we calculated the mean dyadic distance (i.e., the mean inter-individual distance for all possible pairs of group members) as a measure of how tightly packed the group was. At the individual level, we first calculated movement speed (in meters per second) and position relative to the group centroid. Position relative to the centroid is the distance in meters, where a positive y-axis value indicated their distance from the centroid in the direction of group movement, and a positive x-axis value indicated their distance to the right-hand side of the centroid. We calculated two additional measures of each individual’s spatial positions within the group—a surroundedness value (1-CV, where CV is the circular variance of the vectors from group members to the focal individual, [S7]) and their nearest-neighbour distance (i.e., distance in meters to the next closest individual). The former is a reliable measure of centrality (how enclosed an individual is within the group), and the latter provides a measure of how isolated they are. Finally, we summarised these metrics for each ECG window for each focal individual, such that all metrics were summarised into 20 s bursts, with each burst centred on one 6s burst of ECG data. For all metrics, we took the mean over each 20s window as the summary value.

#### Identifying initiation events

To analyse the responses to movement initiation, we extracted successful and unsuccessful movement initiations from the raw GPS data, summarised the co-occurring initiations into events, and identified the level of agreement among initiators. This analysis was conducted using the code from a previously published study on collective decision-making in baboons [S5] and subsequently applied to vulturine guineafowl [S8], see below for more details.

The algorithm we used identifies sequences of initiations from an initial movement of one individual away from another (marked by a change from a minimum to a maximum in their dyadic distance). We defined the initiator as the individual responsible for the majority of the movement that contributed to the increase in dyadic distance. Initiations were considered as successful (a “pull”) if the subsequent return to a minimum dyadic distance was predominately the result of movement by the individual initially left behind (i.e. the ‘potential follower’ followed), and as unsuccessful (an “anchor”) if this was the result of movement by the individual originally identified as the initiator (i.e. the ‘potential follower’ did not follow and the initiator returned). For each initiation attempt, we summarised the number of potential followers as every individual for which there was a maximum within the same sequence of events for the given initiator (these could either become followers or could anchor the initiator after the point at which a maximum dyadic distance was reached, but all were counted as ‘potential followers’).

We also summarised the co-occurring initiations (those in which the “pull” phase of the movement overlapped for a portion of their duration) into initiation events. Then, we identified the level of agreement among the simultaneous initiators using the circular variance (CV) of the unit vectors pointing from the potential follower to each initiator (agreement = 1-CV, where 1 means that all vectors are pointing in the same direction and 0 means that all vectors initiate in opposing directions) [S5]. Note that the number of potential followers was always drawn from the individual-level data (i.e. calculated independently for each initiator) and not aggregated across simultaneous initiators. We used the same code and implementation of this algorithm as [S5], only reducing the minimum change in distance between individuals to 3.5 m (from 5 m in baboons) because guineafowl are substantially smaller and more cohesive. More details of our implementation can be found in [S8].

We then cross-referenced each burst of ECG data with the initiation data, allowing us to identify with each burst whether the individual was initiating (or had recently initiated and either been successful or not), the number of potential followers (i.e. the number of group mates that the individual would ‘pull’ if they were successful in the current initiation attempt) and the extent of the agreement among simultaneous initiators.

#### Statistical analyses

All analyses and data processing steps described here and above were performed in the R environment [S4]. We fitted a total of six LMMs (linear mixed models) using lme4 to test the hypotheses laid out in the main text (three models on change in heart rate relative to resting heart rate, models 1-3, and three models with the same structure but using heart rate variability as response, models 4-6). For all model structures and fixed terms, see tables S1 to S6. Briefly, model 1 tested the effects of movement and spatial positioning on heart rate, model 2 tested the effects of leadership and directional conflict (during initiations) on heart rate, model 3 tested the effect of leadership outcomes and directional conflict (post-initiating), and models 4-6 repeated the same test with heart rate variability as response variable, while maintaining the same model structures models 1-3, respectively. All LMMs included individual IDs and hour of the day as random effects. We only included data when the group was cohesive (∼70% of the data), defined as the mean dyadic distance of individuals being below 25 meters, as it became difficult to determine group and individual-to-group dynamics when the group was more spatially dispersed. Data from two individuals were excluded from the heart rate data because collection peaks could not clearly be identified from their ECG profiles, probably due to poor electrode positioning. Additionally, five birds died, most likely due to an ongoing drought and corresponding increase in predation taking place during the data collection period, but their data until their disappearance was included in all analyses. The data collection resulted in 199,817 data windows for analyses (full sample sizes for all models are given in tables S1 to S6 and in figure captions).

#### Ethics statement

The study was conducted under a research authorisation (KWS/BRP/5001) and capture permit from the Kenyan Wildlife Service (KWS/SCM/5705), the National Commission for Science, Technology and Innovation of Kenya (NACOSTI/P/21/6996), the National Environment Management Authority (NEMA/AGR/68/2017). All the research was done in collaboration with the Ornithology Section of the National Museums of Kenya. Capture and GPS fitting was reviewed by the Max Planck Ethikrat Committee. Implantations were reviewed by the Animal Welfare Officer at the University of Zurich and performed under the supervision of a Kenyan Wildlife Service Vet (Dr. Maureen Kamau). The procedure was reviewed and DZ was approved to perform the surgeries by the Kenya Veterinary Board (KVB/FVS/Voll/6).

## Supplementary Figures

**Fig. S1.**
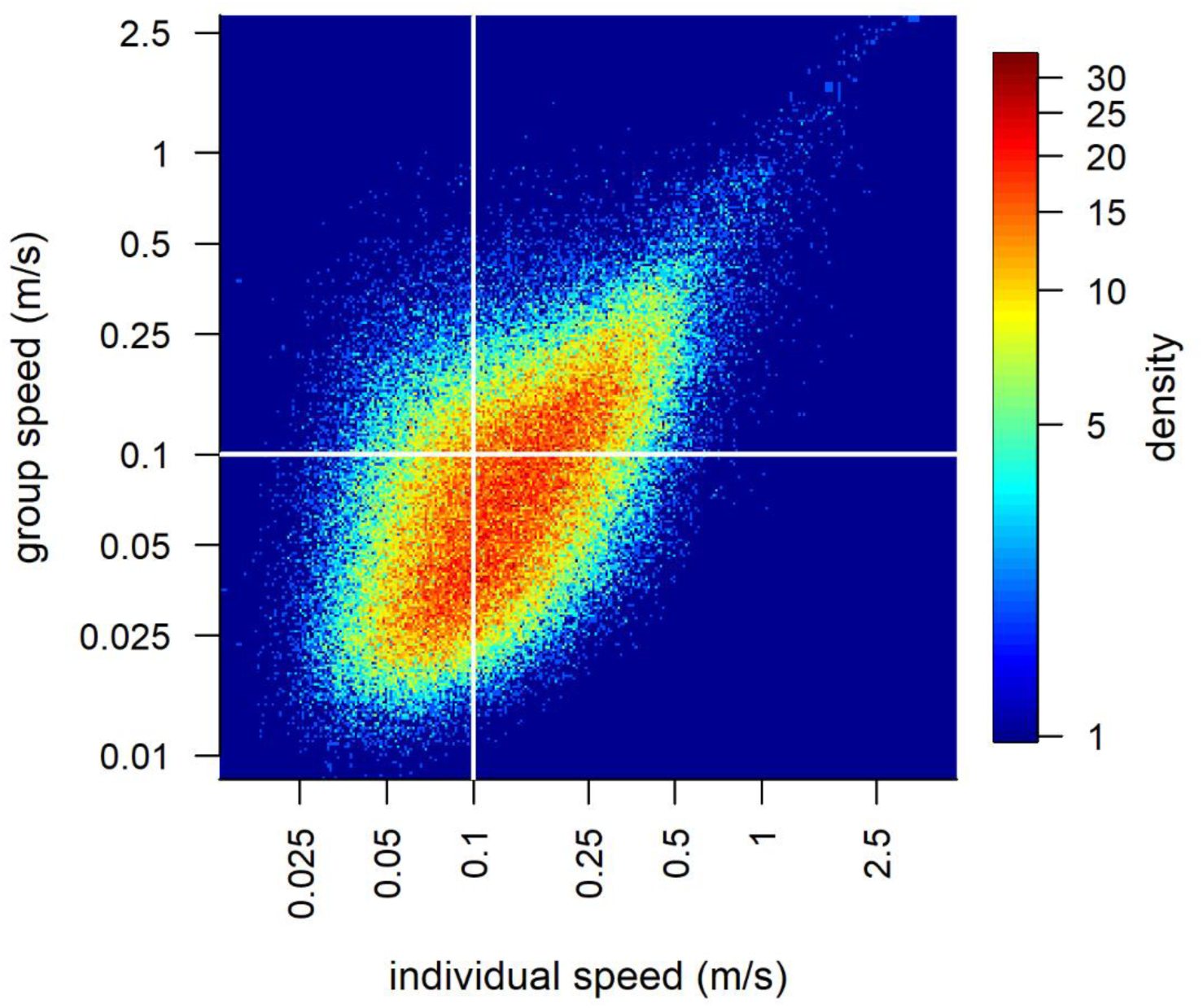
Density plot of individual speeds at different group speeds. The colours represent the density of data points of different individual speeds at different group speeds revealing that individuals mostly move faster than the group (measured as movement of the group centroid).

**Fig. S2.**
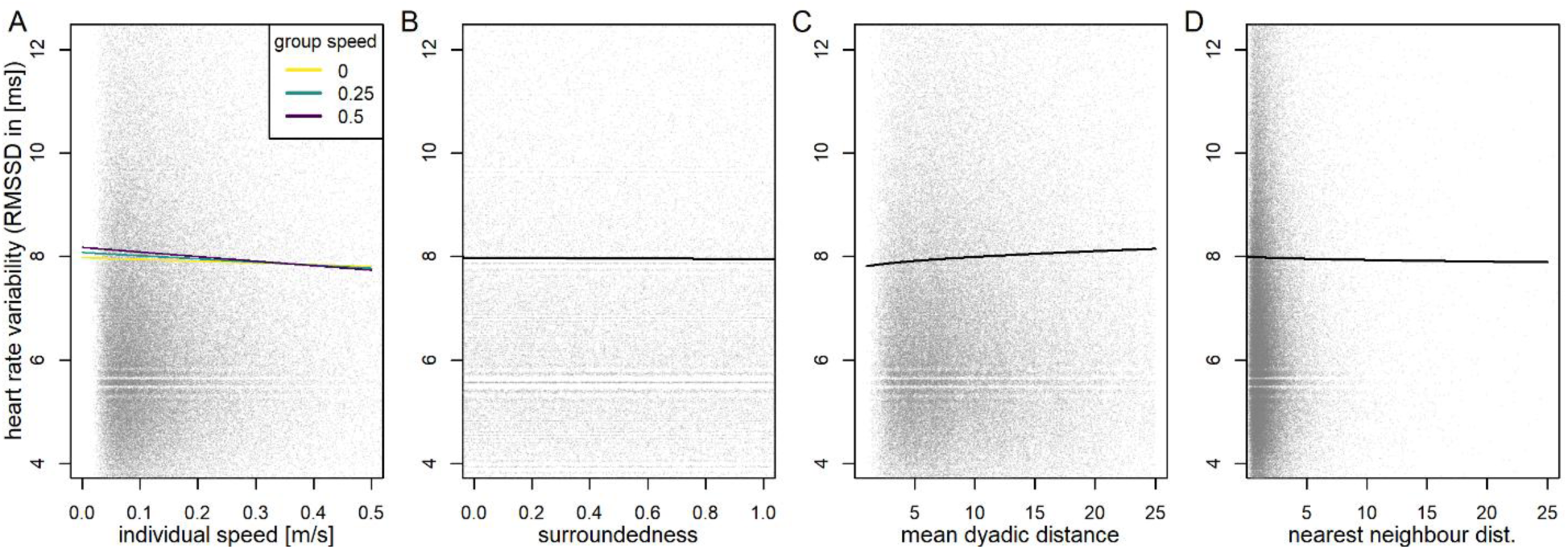
Heart rate variability (HRV) in response to group movement and different within-group positions. (A) Collective movement, (B) being more surrounded, (C) exposure to the front and back of the group, (D) being more proximate to others, and (E) being isolated from other individuals all decrease HRV of vulturine guineafowl. Lines represent the model predictions from predictions from 130,438 ECG data windows (20 s each), back-transformed from the log-scale. Heart rate variability HRV was measured as the RMSSD in ms. All panels are from the same model (model 4, Table S4), and biological average values were used for all values not depicted in each panel. These were: individual speed=0.1 m s^-1^ and group speed=0.1 m s^-1^ (representing slow movements), surroundedness=0.5 (range 0.02–0.99), mean dyadic distance=9.8 (range 1.13–25.0), nearest neighbour distance=2.11 (range 0.06–117.9). Semi-transparent grey points show 80% of the raw data points (lower 5% and upper 15% not shown).

**Fig. S3.**
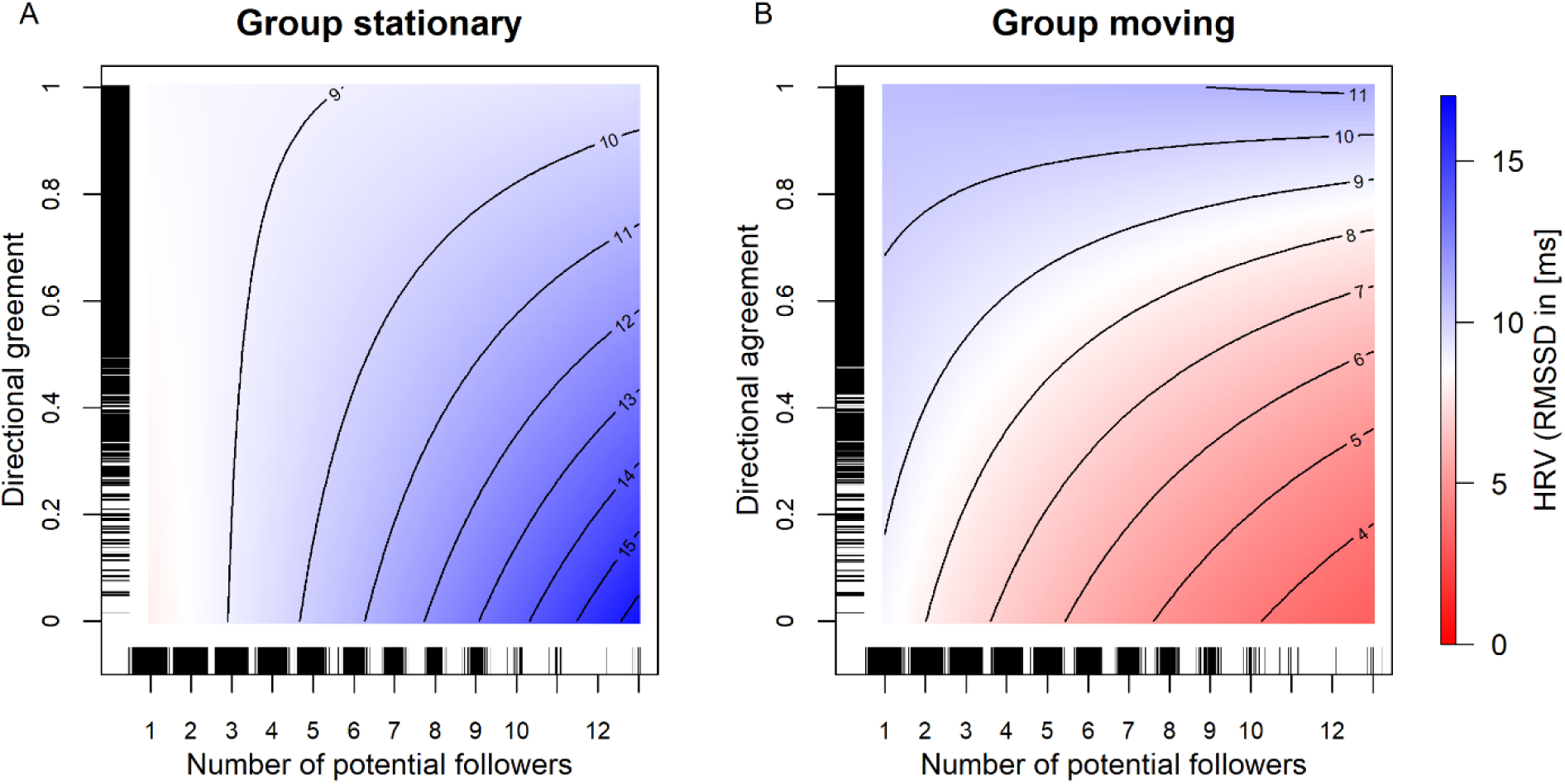
Heart rate variability indicates the greatest signs of stress when there are more potential followers and agreement among initiators is low while the group is moving. Plots show the model predictions from 65,782 ECG data windows (20 s each) back-transformed from the log-scale, in which the focal individual was detected making an initiation movement. The color-coded heart rate variability (HRV) was measured as RMSSD in ms. All panels are from the same model (model 5, Table S5), with group speed set to either 0 m s^-1^ “not moving” (A) or 0.2 m s^-1^ “moving” (B), and all other terms set to 0 as these only had weak effects. Density distributions of raw data points are shown along each of the axes.

**Fig. S4.**
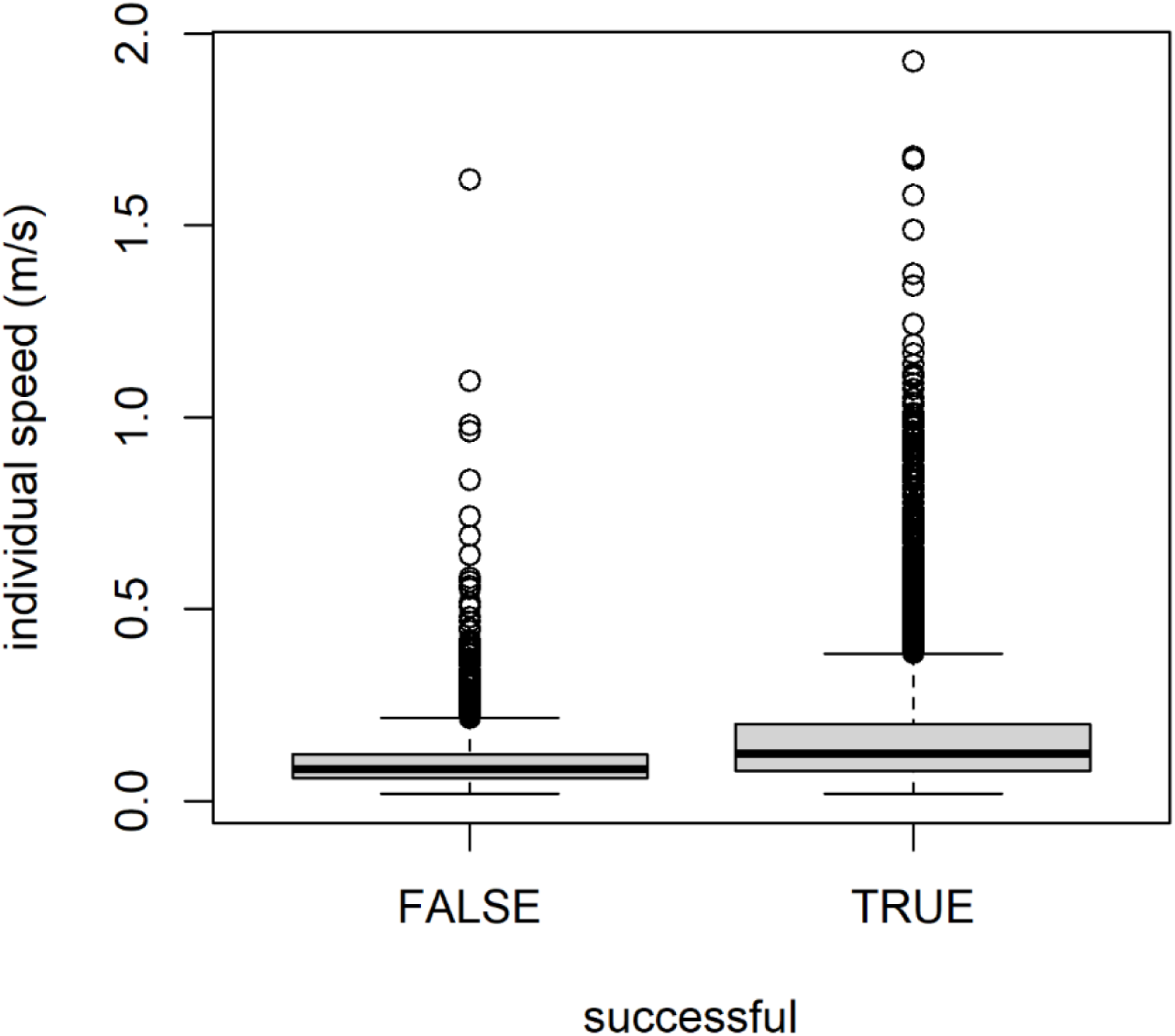
**Average individual speed of movement after an initiation attempt is higher for the successful than the unsuccessful initiators.**

**Fig. S5.**
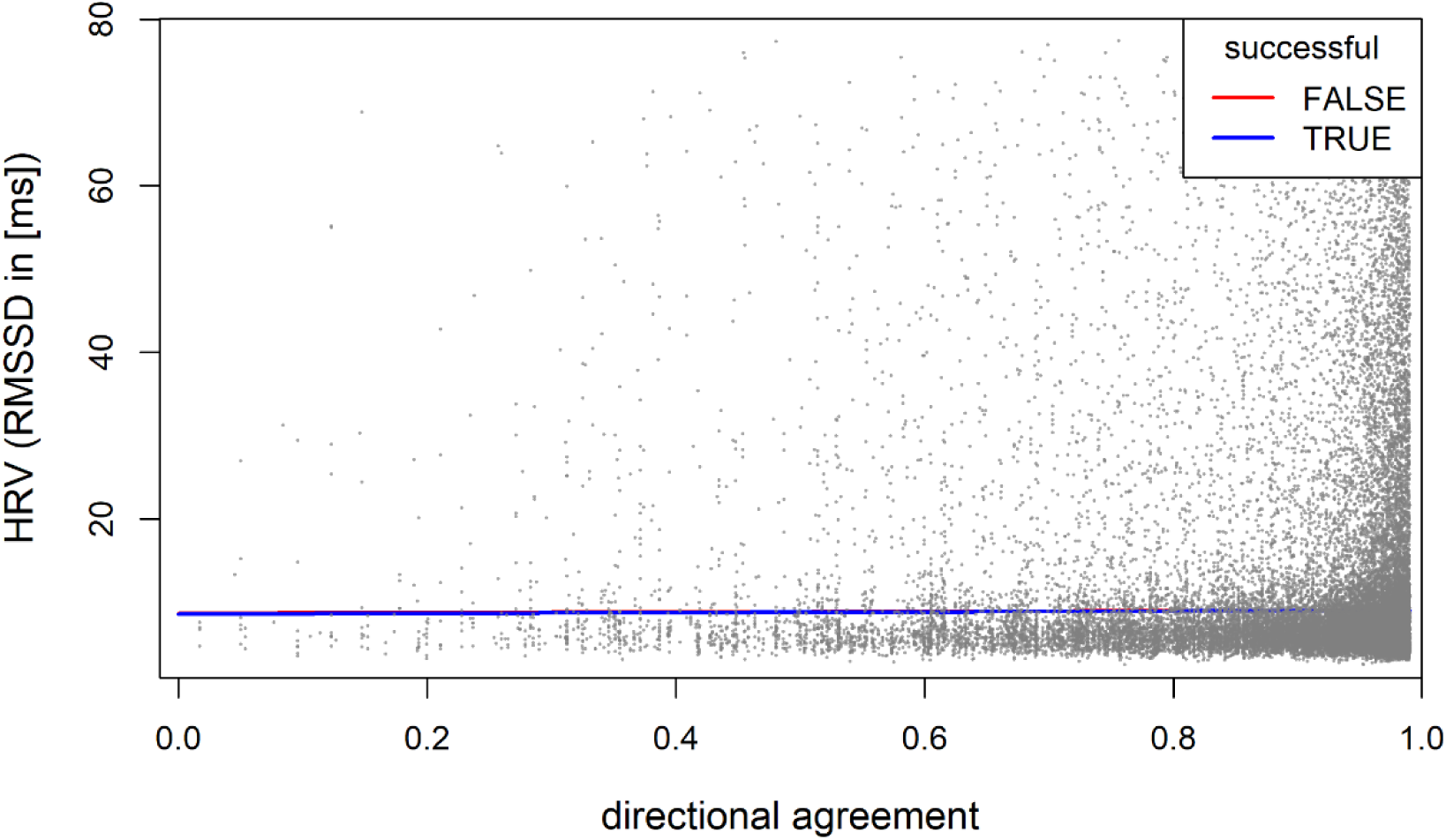
Heart rate variability in response to group movement and different within-group positions. Lines show the model predictions back-transformed from the log-scale from 30,083 ECG data windows (20 s each) where an individual was either successful (TRUE) or unsuccessful (FALSE) in initiating. Heart rate variability (HRV) was measured as the RMSSD in ms. The fits are from model 6 (table 6), with all other terms set to 0 as these only scaled the results up or down along the y-axis. Rug plots along the axes show the distribution of 95% of the raw data (not including lower and upper 2.5%).

---

**Table S1.**
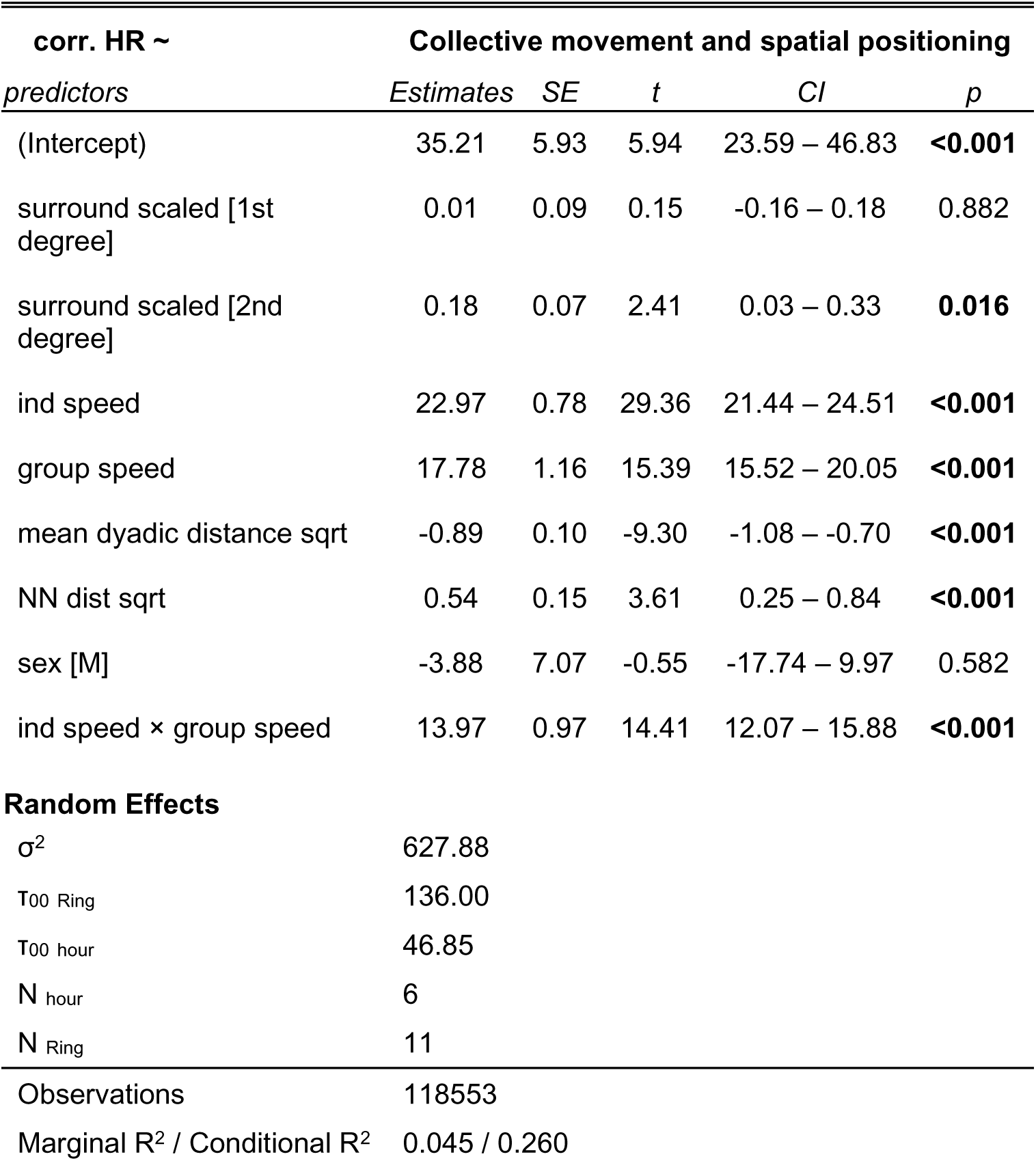
Model 1 results – Effect of collective movement and spatial positioning on heart rate.

**Table S2.**
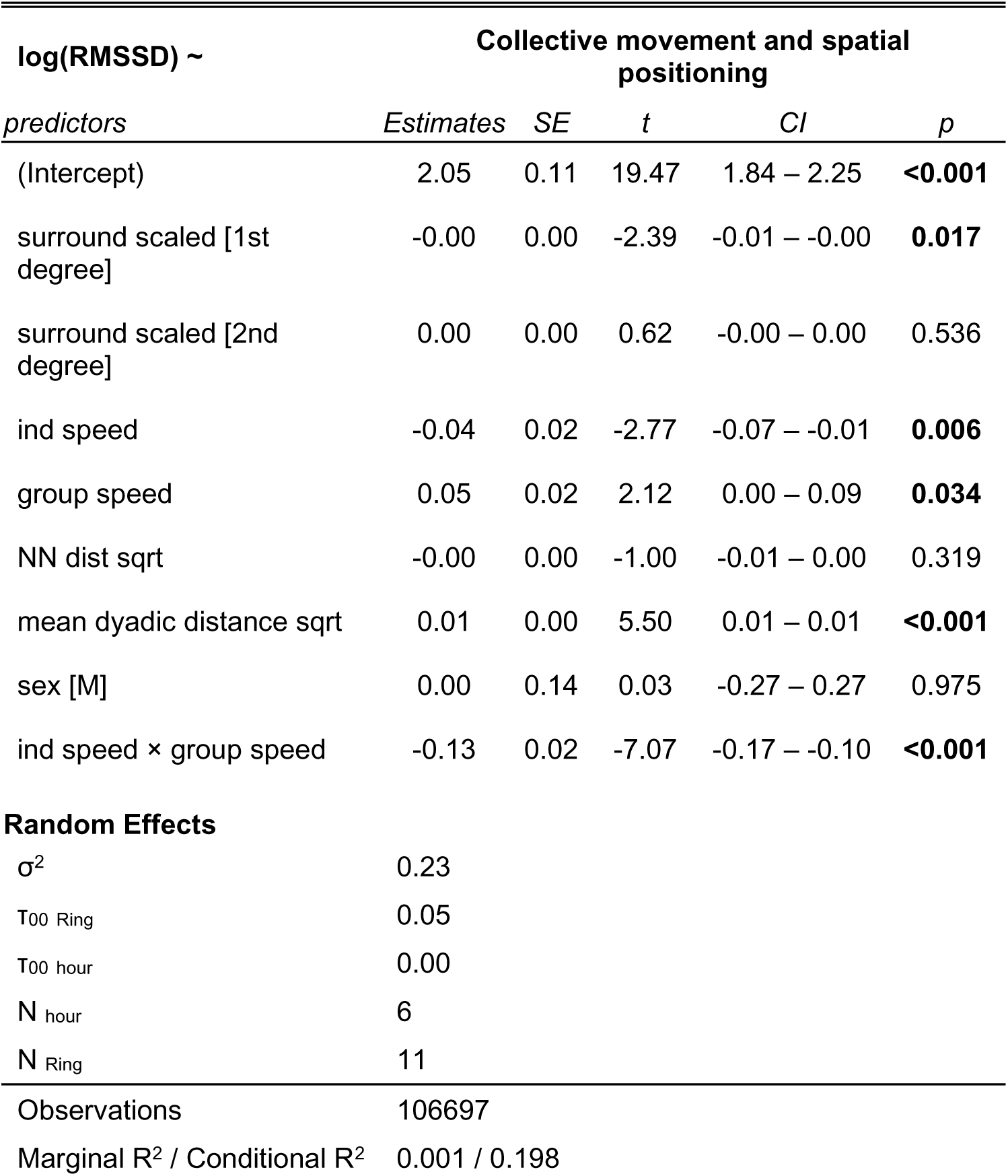
Model 4 results – Effect of collective movement and spatial positioning on heart rate variability (measured as log(RMSSD) in ms).

**Table S3.**
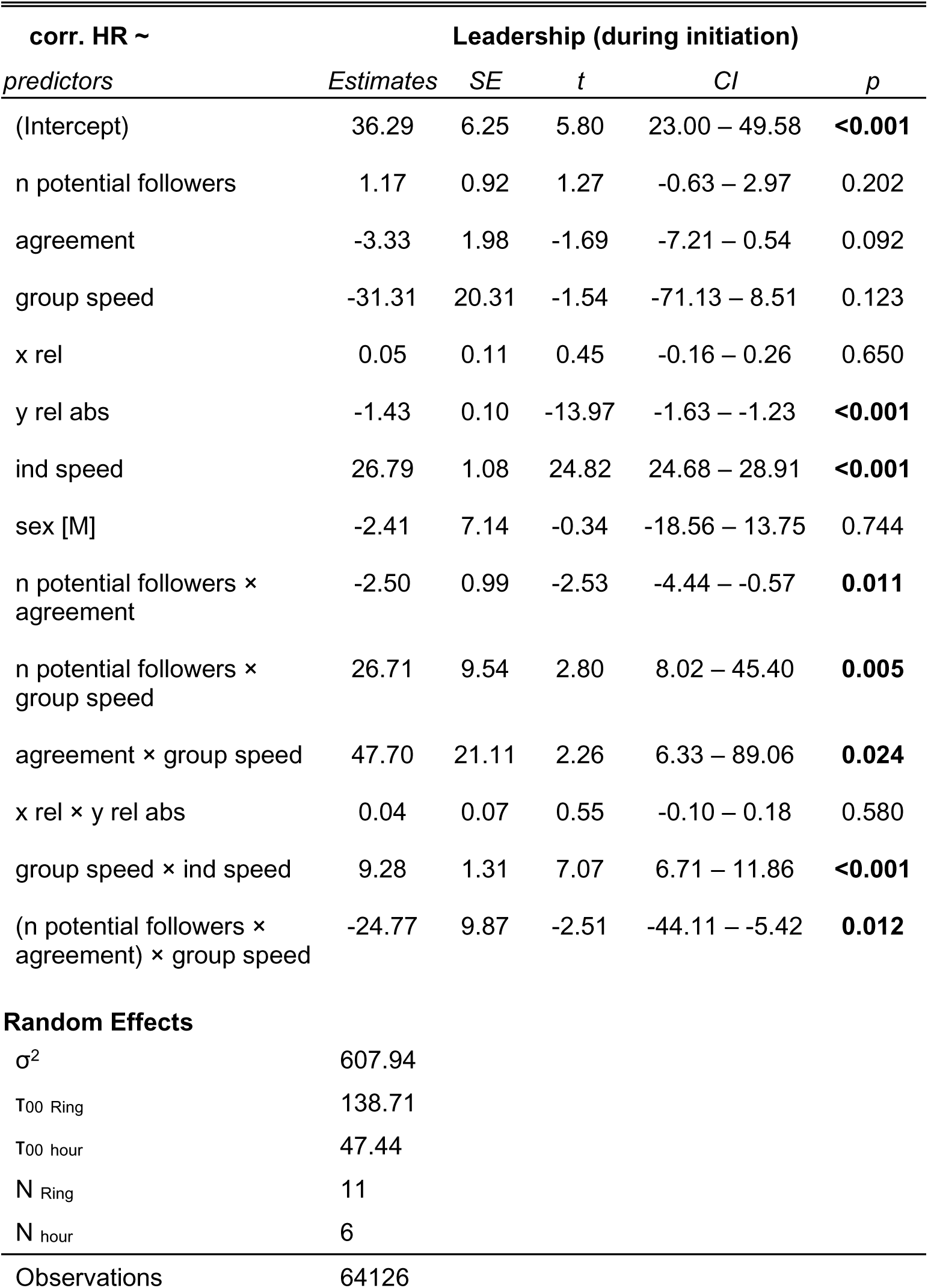
Model 2 results – Effects of leadership and directional conflict (during initiations) on heart rate.

**Table S4.**
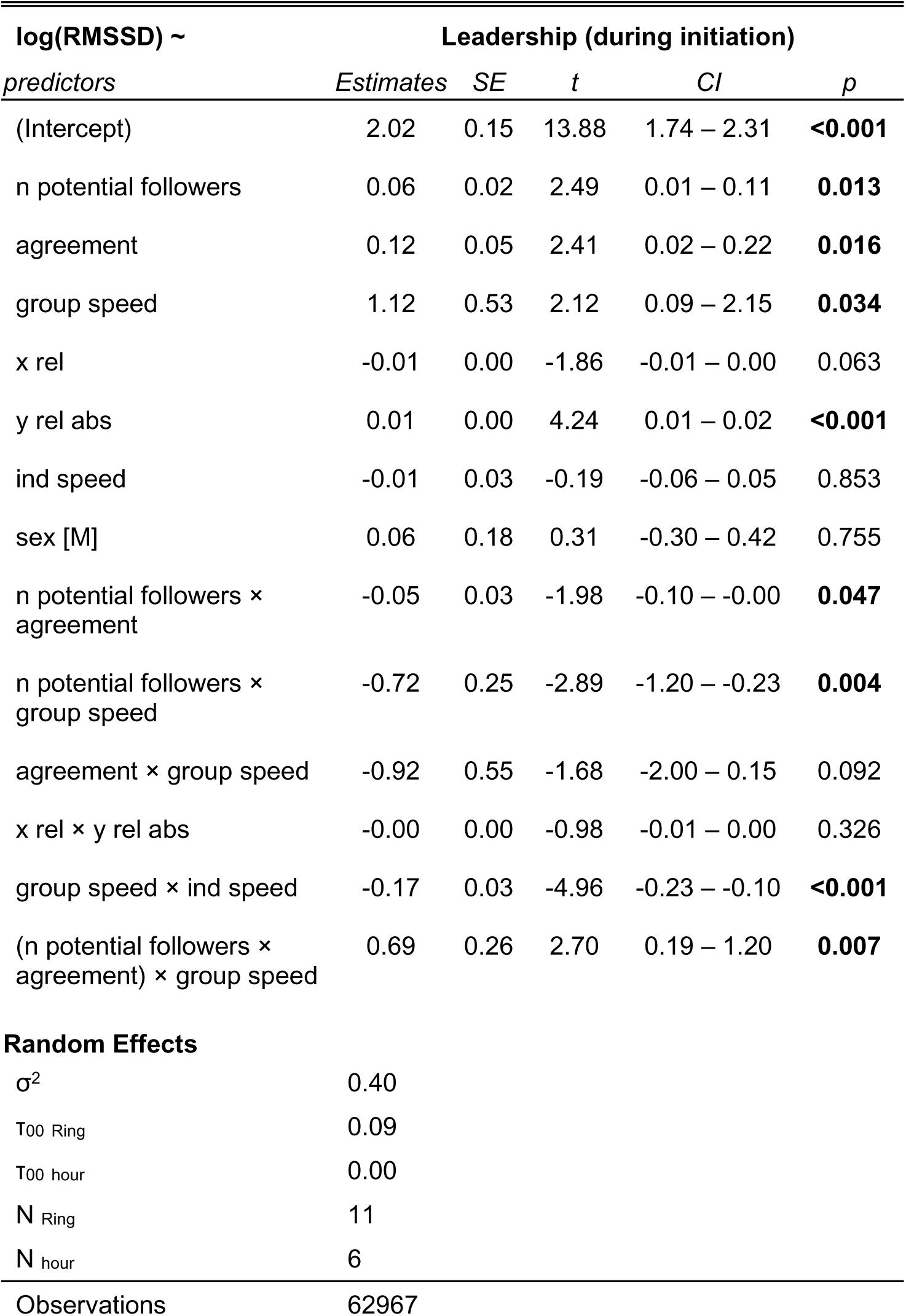
Model 5 results – Effect of leadership and directional disagreement (during initiation) on heart rate variability (measured as log(RMSSD) in ms).

**Table S5.**
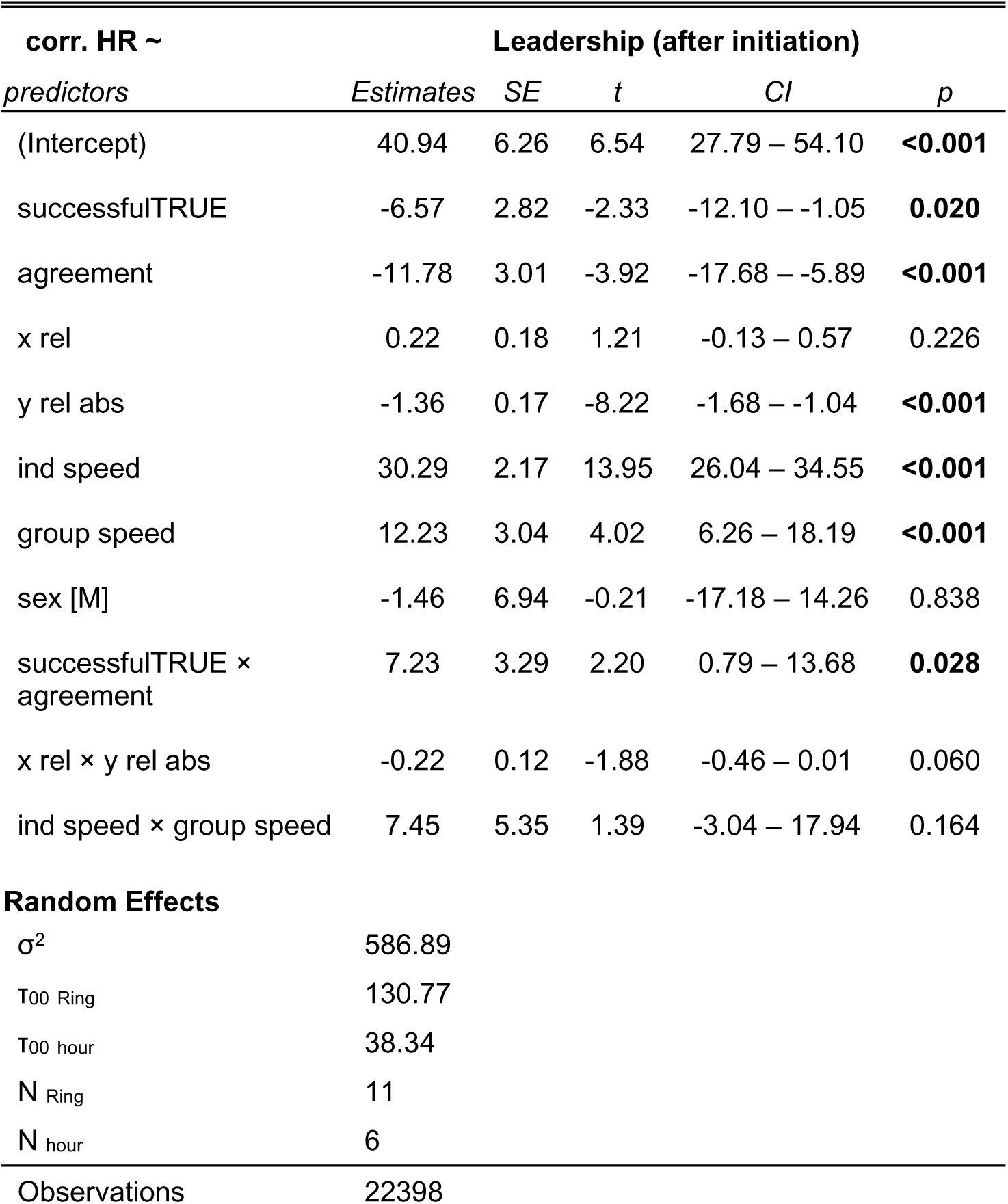
Model 3 results – Effects of leadership outcomes and directional disagreement (after initiation) on heart rate.

**Table S6.**
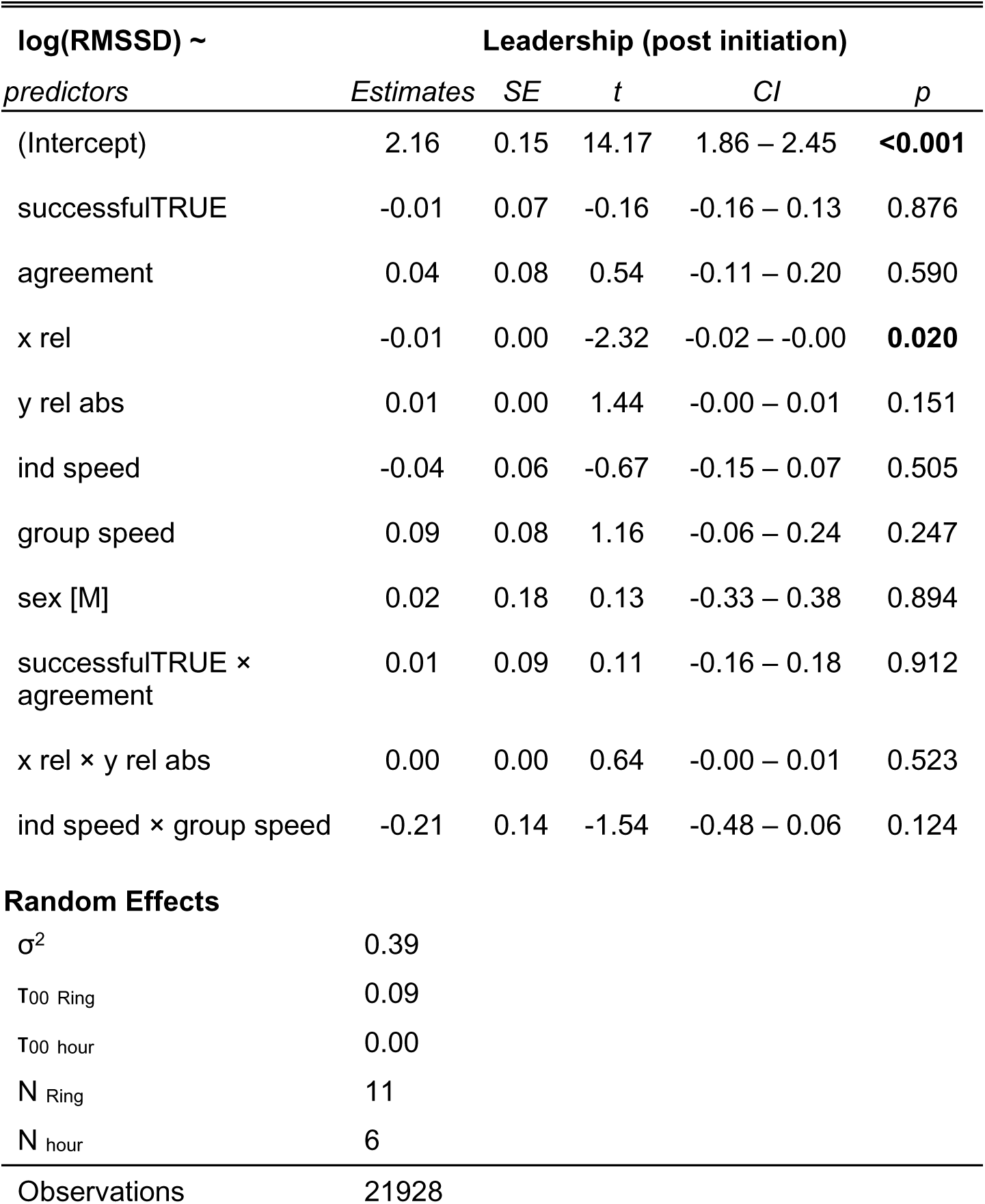
Model 6 results – Effects of leadership outcomes and directional disagreement (after initiation) on heart rate variability (measured as log(RMSSD) in ms).

**Movie S1. Example of collective movement by a group of vulturine guineafowl**. This demonstrates the high cohesion among group members and the dynamic spatial positioning of individuals.

