## Supplementary Methods and Materials for "The physiological costs of leadership in collective movements"

**Supplementary Figures**

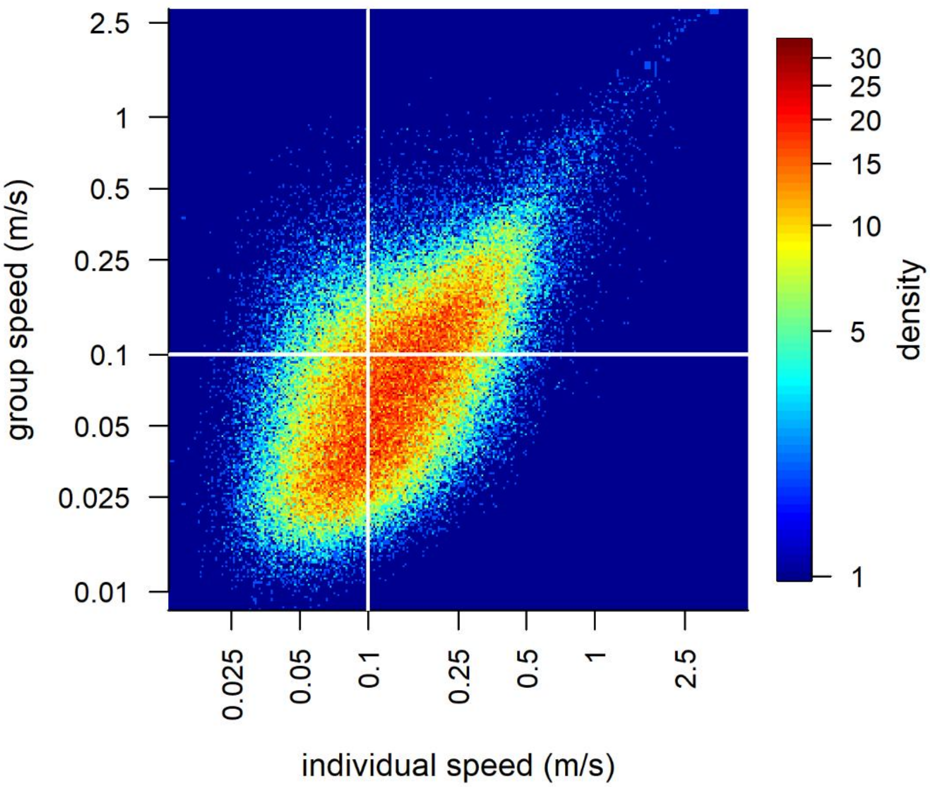


**Fig. S1. Density plot of individual speeds at different group speeds.**

The colours represent the density of data points of different individual speeds at different group speeds revealing that individuals mostly move faster than the group (measured as movement of the group centroid).

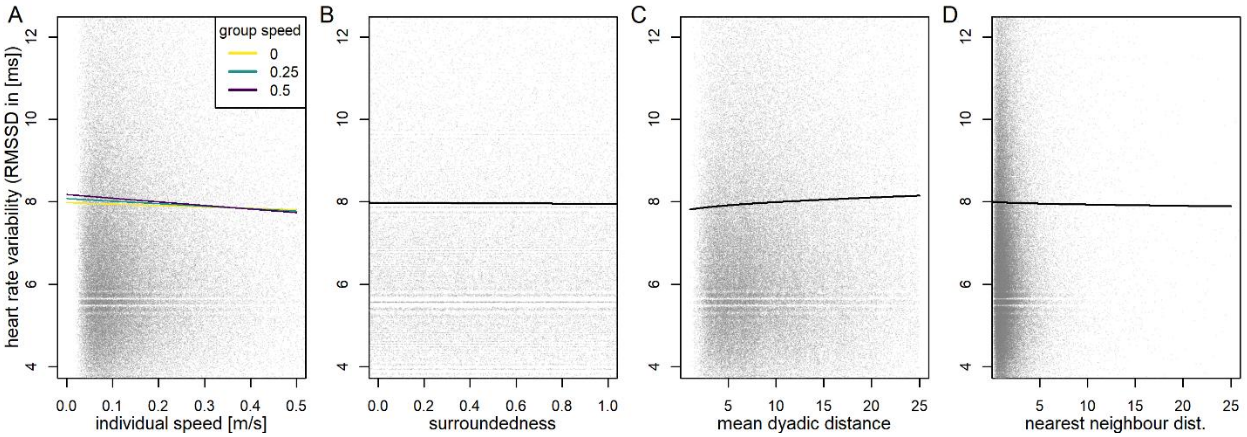


**Fig. S2. Heart rate variability (HRV) in response to group movement and different within-group positions.**

(A) Collective movement, (B) being more surrounded, (C) exposure to the front and back of the group, (D) being more proximate to others, and (E) being isolated from other individuals all decrease HRV of vulturine guineafowl. Lines represent the model predictions from predictions from 130,438 ECG data windows (20 s each), back-transformed from the log-scale. Heart rate variability HRV was measured as the RMSSD in ms. All panels are from the same model (model 4, table S4), and biological average values were used for all values not depicted in each panel. These were: individual speed=0.1 m $s^{-1}$ and group speed=0.1 m $s^{-1}$ (representing slow movements), surroundedness=0.5 (range 0.02–0.99), mean dyadic distance=9.8 (range 1.13–25.0), nearest neighbour distance=2.11 (range 0.06–117.9). Semi-transparent grey points show 80% of the raw data points (lower 5% and upper 15% not shown).

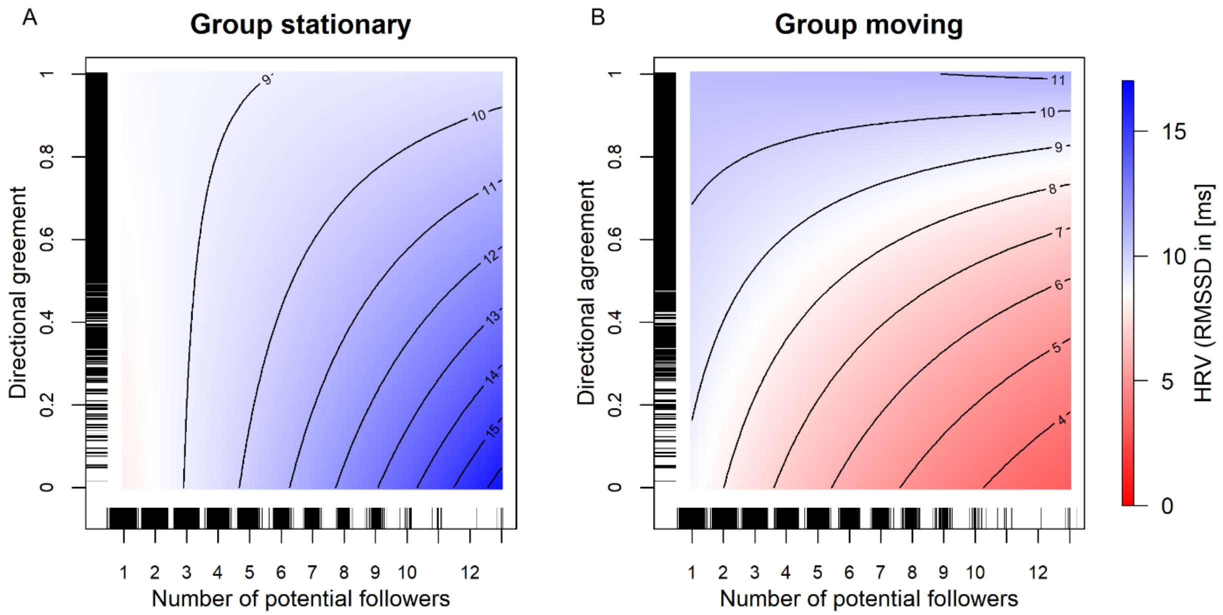


**Fig. S3. Heart rate variability indicates the greatest signs of stress when there are more potential followers and agreement among initiators is low while the group is moving.**

Plots show the model predictions from 65,782 ECG data windows (20 s each) back-transformed from the log-scale, in which the focal individual was detected making an initiation movement. The color-coded heart rate variability (HRV) was measured as RMSSD in ms. All panels are from the same model (model 5, table S5), with group speed set to either 0 m $s^{-1}$ "not moving" (A) or 0.2 m $s^{-1}$ "moving" (B), and all other terms set to 0 as these only had weak effects. Density distributions of raw data points are shown along each of the axes.

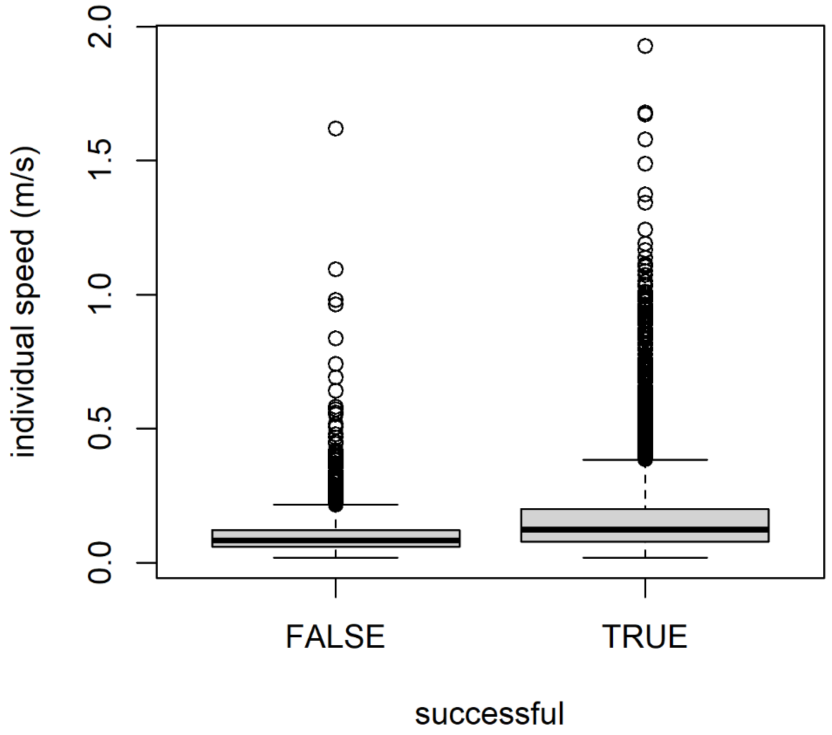


**Fig. S4. Average individual speed of movement after an initiation attempt is higher for the successful than the unsuccessful initiators.**

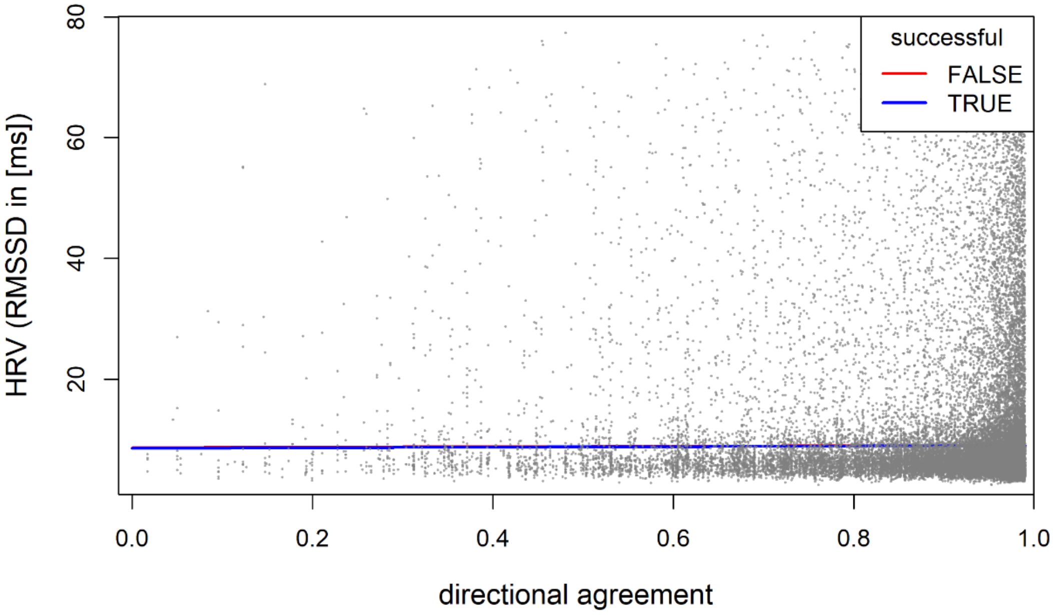


**Fig. S5. Heart rate variability in response to group movement and different within-group positions.**

Lines show the model predictions back-transformed from the log-scale from 30,083 ECG data windows (20 s each) where an individual was either successful (TRUE) or unsuccessful (FALSE) in initiating. Heart rate variability (HRV) was measured as the RMSSD in ms. The fits are from model 6 (table 6), with all other terms set to 0 as these only scaled the results up or down along the y-axis. Rug plots along the axes show the distribution of 95% of the raw data (not including lower and upper 2.5%).

**Table S1. Model 1 results – Effect of collective movement and spatial positioning on heart rate.**

| **corr. HR ~** | **Collective movement and spatial positioning** | | | | |
|---|---|---|---|---|---|
| *predictors* | *Estimates* | *SE* | *t* | *CI* | *p* |
| (Intercept) | 35.21 | 5.93 | 5.94 | 23.59 – 46.83 | **<0.001** |
| surround scaled [1st degree] | 0.01 | 0.09 | 0.15 | -0.16 – 0.18 | 0.882 |
| surround scaled [2nd degree] | 0.18 | 0.07 | 2.41 | 0.03 – 0.33 | **0.016** |
| ind speed | 22.97 | 0.78 | 29.36 | 21.44 – 24.51 | **<0.001** |
| group speed | 17.78 | 1.16 | 15.39 | 15.52 – 20.05 | **<0.001** |
| mean dyadic distance sqrt | -0.89 | 0.10 | -9.30 | -1.08 – -0.70 | **<0.001** |
| NN dist sqrt | 0.54 | 0.15 | 3.61 | 0.25 – 0.84 | **<0.001** |
| sex [M] | -3.88 | 7.07 | -0.55 | -17.74 – 9.97 | 0.582 |
| ind speed × group speed | 13.97 | 0.97 | 14.41 | 12.07 – 15.88 | **<0.001** |
| **Random Effects** | | | | | |
| $\sigma^2$ | 627.88 | | | | |
| $\tau_{00\ Ring}$ | 136.00 | | | | |
| $\tau_{00\ hour}$ | 46.85 | | | | |
| $N_{hour}$ | 6 | | | | |
| $N_{Ring}$ | 11 | | | | |
| Observations | 118553 | | | | |
| Marginal $R^2$ / Conditional $R^2$ | 0.045 / 0.260 | | | | |

**Table S2. Model 4 results – Effect of collective movement and spatial positioning on heart rate variability (measured as log(RMSSD) in ms).**

| **log(RMSSD) ~** | **Collective movement and spatial positioning** | | | | |
|---|---|---|---|---|---|
| *predictors* | *Estimates* | *SE* | *t* | *CI* | *p* |
| (Intercept) | 2.05 | 0.11 | 19.47 | 1.84 – 2.25 | **<0.001** |
| surround scaled [1st degree] | -0.00 | 0.00 | -2.39 | -0.01 – -0.00 | **0.017** |
| surround scaled [2nd degree] | 0.00 | 0.00 | 0.62 | -0.00 – 0.00 | 0.536 |
| ind speed | -0.04 | 0.02 | -2.77 | -0.07 – -0.01 | **0.006** |
| group speed | 0.05 | 0.02 | 2.12 | 0.00 – 0.09 | **0.034** |
| NN dist sqrt | -0.00 | 0.00 | -1.00 | -0.01 – 0.00 | 0.319 |
| mean dyadic distance sqrt | 0.01 | 0.00 | 5.50 | 0.01 – 0.01 | **<0.001** |
| sex [M] | 0.00 | 0.14 | 0.03 | -0.27 – 0.27 | 0.975 |
| ind speed × group speed | -0.13 | 0.02 | -7.07 | -0.17 – -0.10 | **<0.001** |
| **Random Effects** | | | | | |
| $\sigma^2$ | 0.23 | | | | |
| $\tau_{00\ Ring}$ | 0.05 | | | | |
| $\tau_{00\ hour}$ | 0.00 | | | | |
| $N_{hour}$ | 6 | | | | |
| $N_{Ring}$ | 11 | | | | |
| Observations | 106697 | | | | |
| Marginal $R^2$ / Conditional $R^2$ | 0.001 / 0.198 | | | | |

**Table S3. Model 2 results – Effects of leadership and directional conflict (during initiations) on heart rate.**

| **corr. HR ~** | **Leadership (during initiation)** | | | | |
|---|---|---|---|---|---|
| *predictors* | *Estimates* | *SE* | *t* | *CI* | *p* |
| (Intercept) | 36.29 | 6.25 | 5.80 | 23.00 – 49.58 | **<0.001** |
| n potential followers | 1.17 | 0.92 | 1.27 | -0.63 – 2.97 | 0.202 |
| agreement | -3.33 | 1.98 | -1.69 | -7.21 – 0.54 | 0.092 |
| group speed | -31.31 | 20.31 | -1.54 | -71.13 – 8.51 | 0.123 |
| x rel | 0.05 | 0.11 | 0.45 | -0.16 – 0.26 | 0.650 |
| y rel abs | -1.43 | 0.10 | -13.97 | -1.63 – -1.23 | **<0.001** |
| ind speed | 26.79 | 1.08 | 24.82 | 24.68 – 28.91 | **<0.001** |
| sex [M] | -2.41 | 7.14 | -0.34 | -18.56 – 13.75 | 0.744 |
| n potential followers × agreement | -2.50 | 0.99 | -2.53 | -4.44 – -0.57 | **0.011** |
| n potential followers × group speed | 26.71 | 9.54 | 2.80 | 8.02 – 45.40 | **0.005** |
| agreement × group speed | 47.70 | 21.11 | 2.26 | 6.33 – 89.06 | **0.024** |
| x rel × y rel abs | 0.04 | 0.07 | 0.55 | -0.10 – 0.18 | 0.580 |
| group speed × ind speed | 9.28 | 1.31 | 7.07 | 6.71 – 11.86 | **<0.001** |
| (n potential followers × agreement) × group speed | -24.77 | 9.87 | -2.51 | -44.11 – -5.42 | **0.012** |
| **Random Effects** | | | | | |
| $\sigma^2$ | 607.94 | | | | |
| $\tau_{00\ Ring}$ | 138.71 | | | | |
| $\tau_{00\ hour}$ | 47.44 | | | | |
| $N_{Ring}$ | 11 | | | | |
| $N_{hour}$ | 6 | | | | |
| Observations | 64126 | | | | |

**Table S4. Model 5 results – Effect of leadership and directional disagreement (during initiation) on heart rate variability (measured as log(RMSSD) in ms).**

| **log(RMSSD) ~** | **Leadership (during initiation)** | | | | |
|---|---|---|---|---|---|
| *predictors* | *Estimates* | *SE* | *t* | *CI* | *p* |
| (Intercept) | 2.02 | 0.15 | 13.88 | 1.74 – 2.31 | **<0.001** |
| n potential followers | 0.06 | 0.02 | 2.49 | 0.01 – 0.11 | **0.013** |
| agreement | 0.12 | 0.05 | 2.41 | 0.02 – 0.22 | **0.016** |
| group speed | 1.12 | 0.53 | 2.12 | 0.09 – 2.15 | **0.034** |
| x rel | -0.01 | 0.00 | -1.86 | -0.01 – 0.00 | 0.063 |
| y rel abs | 0.01 | 0.00 | 4.24 | 0.01 – 0.02 | **<0.001** |
| ind speed | -0.01 | 0.03 | -0.19 | -0.06 – 0.05 | 0.853 |
| sex [M] | 0.06 | 0.18 | 0.31 | -0.30 – 0.42 | 0.755 |
| n potential followers × agreement | -0.05 | 0.03 | -1.98 | -0.10 – -0.00 | **0.047** |
| n potential followers × group speed | -0.72 | 0.25 | -2.89 | -1.20 – -0.23 | **0.004** |
| agreement × group speed | -0.92 | 0.55 | -1.68 | -2.00 – 0.15 | 0.092 |
| x rel × y rel abs | -0.00 | 0.00 | -0.98 | -0.01 – 0.00 | 0.326 |
| group speed × ind speed | -0.17 | 0.03 | -4.96 | -0.23 – -0.10 | **<0.001** |
| (n potential followers × agreement) × group speed | 0.69 | 0.26 | 2.70 | 0.19 – 1.20 | **0.007** |
| **Random Effects** | | | | | |
| $\sigma^2$ | 0.40 | | | | |
| $\tau_{00\ Ring}$ | 0.09 | | | | |
| $\tau_{00\ hour}$ | 0.00 | | | | |
| $N_{Ring}$ | 11 | | | | |
| $N_{hour}$ | 6 | | | | |
| Observations | 62967 | | | | |

**Table S5. Model 3 results – Effects of leadership outcomes and directional disagreement (after initiation) on heart rate.**

| **corr. HR ~** | **Leadership (after initiation)** | | | | |
|---|---|---|---|---|---|
| *predictors* | *Estimates* | *SE* | *t* | *CI* | *p* |
| (Intercept) | 40.94 | 6.26 | 6.54 | 27.79 – 54.10 | **<0.001** |
| successfulTRUE | -6.57 | 2.82 | -2.33 | -12.10 – -1.05 | **0.020** |
| agreement | -11.78 | 3.01 | -3.92 | -17.68 – -5.89 | **<0.001** |
| x rel | 0.22 | 0.18 | 1.21 | -0.13 – 0.57 | 0.226 |
| y rel abs | -1.36 | 0.17 | -8.22 | -1.68 – -1.04 | **<0.001** |
| ind speed | 30.29 | 2.17 | 13.95 | 26.04 – 34.55 | **<0.001** |
| group speed | 12.23 | 3.04 | 4.02 | 6.26 – 18.19 | **<0.001** |
| sex [M] | -1.46 | 6.94 | -0.21 | -17.18 – 14.26 | 0.838 |
| successfulTRUE × agreement | 7.23 | 3.29 | 2.20 | 0.79 – 13.68 | **0.028** |
| x rel × y rel abs | -0.22 | 0.12 | -1.88 | -0.46 – 0.01 | 0.060 |
| ind speed × group speed | 7.45 | 5.35 | 1.39 | -3.04 – 17.94 | 0.164 |
| **Random Effects** | | | | | |
| $\sigma^2$ | 586.89 | | | | |
| $\tau_{00\ Ring}$ | 130.77 | | | | |
| $\tau_{00\ hour}$ | 38.34 | | | | |
| $N_{Ring}$ | 11 | | | | |
| $N_{hour}$ | 6 | | | | |
| Observations | 22398 | | | | |

**Table S6. Model 6 results – Effects of leadership outcomes and directional disagreement (after initiation) on heart rate variability (measured as log(RMSSD) in ms).**

| **log(RMSSD) ~** | **Leadership (post initiation)** | | | | |
|---|---|---|---|---|---|
| *predictors* | *Estimates* | *SE* | *t* | *CI* | *p* |
| (Intercept) | 2.16 | 0.15 | 14.17 | 1.86 – 2.45 | **<0.001** |
| successfulTRUE | -0.01 | 0.07 | -0.16 | -0.16 – 0.13 | 0.876 |
| agreement | 0.04 | 0.08 | 0.54 | -0.11 – 0.20 | 0.590 |
| x rel | -0.01 | 0.00 | -2.32 | -0.02 – -0.00 | **0.020** |
| y rel abs | 0.01 | 0.00 | 1.44 | -0.00 – 0.01 | 0.151 |
| ind speed | -0.04 | 0.06 | -0.67 | -0.15 – 0.07 | 0.505 |
| group speed | 0.09 | 0.08 | 1.16 | -0.06 – 0.24 | 0.247 |
| sex [M] | 0.02 | 0.18 | 0.13 | -0.33 – 0.38 | 0.894 |
| successfulTRUE × agreement | 0.01 | 0.09 | 0.11 | -0.16 – 0.18 | 0.912 |
| x rel × y rel abs | 0.00 | 0.00 | 0.64 | -0.00 – 0.01 | 0.523 |
| ind speed × group speed | -0.21 | 0.14 | -1.54 | -0.48 – 0.06 | 0.124 |
| **Random Effects** | | | | | |
| $\sigma^2$ | 0.39 | | | | |
| $\tau_{00\ Ring}$ | 0.09 | | | | |
| $\tau_{00\ hour}$ | 0.00 | | | | |
| $N_{Ring}$ | 11 | | | | |
| $N_{hour}$ | 6 | | | | |
| Observations | 21928 | | | | |

**Movie S1. Example of collective movement by a group of vulturine guineafowl**. This demonstrates the high cohesion among group members and the dynamic spatial positioning of individuals.
